# Microglia ferroptosis is prevalent in neurodegenerative disease and regulated by SEC24B

**DOI:** 10.1101/2021.11.02.466996

**Authors:** Sean K. Ryan, Matija Zelic, Yingnan Han, Erin Teeple, Luoman Chen, Mahdiar Sadeghi, Srinivas Shankara, Lilu Guo, Cong Li, Fabrizio Pontarelli, Elizabeth H. Jensen, Dinesh Kumar, Mindy Zhang, Joseph Gans, Bailin Zhang, Jonathan Proto, Jacqueline Saleh, James C. Dodge, Deepak Rajpal, Dimitry Ofengeim, Timothy R. Hammond

## Abstract

Iron dysregulation has been implicated in multiple neurodegenerative diseases, including Parkinson’s Disease (PD), Amyotrophic Lateral Sclerosis (ALS), and Multiple Sclerosis (MS). One prominent feature of affected brain regions are iron-loaded microglia, but how iron overload influences microglia physiology and disease response is poorly understood. Here we show that microglia are highly susceptible to ferroptosis, an iron-dependent form of cell death. In a tri-culture of human iPSC-derived neurons, astrocytes, and microglia, under ferroptosis-inducing conditions, microglia undergo a drastic shift in cell state, with increased ferritin levels, disrupted glutathione homeostasis, and altered cytokine signaling. Similar ferroptosis-associated signature (FAS) microglia were uncovered in PD, and the signature was also found in a large cohort of PD patient blood samples, raising the possibility that ferroptosis can be identified clinically. We performed a genome-wide CRISPR screen which revealed a novel regulator of ferroptosis, the vesicle trafficking gene SEC24B. A small molecule screen also nominated several candidates which blocked ferroptosis, some of which are already in clinical use. These data suggest that ferroptosis sits at the interface of cell death and inflammation, and inhibition of this process in microglia and other brain cells may provide new ways for treating neurodegenerative disease.

## Introduction

Iron is important for redox-based metabolic activities and is the most abundant transition metal in the brain [1]. Disrupted iron homeostasis has been implicated in neurodegeneration [2] and iron accumulation has been correlated with disease progression in several neurodegenerative disorders including Parkinson’s disease (PD), Amyotrophic Lateral Sclerosis (ALS), and Friedreich’s Ataxia [3–5]. While all cell types in the brain can store iron, microglia have one of the highest storage capacities and are prone to iron accumulation in disease [1, 6–10]. Additionally, a subpopulation of iron-laden microglia with a unique transcriptomic signature has been discovered in the rim of progressive multiple sclerosis (MS) lesions [11, 12], raising questions about how these cells participate in pathological progression in MS as well as other neuroinflammatory and neurodegenerative disorders

One mechanism that has not been explored in detail is whether microglia function is altered in response to iron dysregulation and whether these cells are susceptible to a novel iron-dependent form of cell death called ferroptosis [13]. Ferroptosis is distinct from other forms of cell death like apoptosis and necroptosis and is driven by iron-dependent phospholipid peroxidation [13]. Ferroptosis has been implicated in multiple neurodegenerative disorders including PD and mutations in the iron storage gene FTL cause a rare form of Parkinsonism [14]. Interestingly, disease-relevant subtypes of neurons including motor neurons and dopaminergic neurons seem especially susceptible to ferroptosis [15, 16], but the role in glia is largely unexplored. Previous studies have shown that primary and immortalized mouse microglia can also undergo ferroptosis in mono-cultures [17], but the effect of microglial iron accumulation in cellular function or neurodegeneration is not fully understood.

To understand the role of iron accumulation in human microglia and to model the complex interactions between neurons and glia in a disease-relevant system, we developed a human induced pluripotent stem cell (hiPSC)-derived tri-culture system that contained microglia, neurons, and astrocytes [18]. We found that microglia had the strongest transcriptional response to iron dysregulation among the three cell types. Using single cell RNA sequencing (scRNAseq), we identify a subset of microglia with a distinct ferroptosis-associated signature (FAS). We also found enrichment of the microglia iron dysregulation/FAS in the spinal cord of ALS patients, as well as, in blood from two large PD patient cohorts and in microglia from single nuclei RNAseq (snRNAseq) from PD patient midbrain samples. To understand how iron-dependent signaling is regulated in microglia, we performed a genome-wide CRISPR screen and identified a network that regulates ferroptosis in microglia. Interestingly, we identified key genes that regulate this form of cell death in microglia including *ACSL4* and a novel ferroptosis susceptibility gene *SEC24B*. Finally, we performed a small molecule screen to identify inhibitors of this process in microglia and showed that pharmacological modulation may be a viable strategy to mitigate ferroptosis in neurodegenerative disease. These findings point to an important role for ferroptosis in microglia that may interplay with neuronal ferroptosis.

## Results

### Induction of ferroptosis in human iPSC tri-culture reveals prominent microglia ferroptotic signature

To better understand the role of iron signaling and ferroptosis in the brain, we made a human iPSC-derived tri-culture of neurons, astrocytes, and microglia (Fig. 1A) [18, 19]. This system can be used for acute and long-term studies and consists of approximately 15% microglia, 25% neurons, and 60% astrocytes as determined by FACS and immunocytochemistry (Fig. 1B and S1A). All cell types were well integrated and form a complex network within two weeks (Fig. 1C). To study the role of iron overload and whether these cultures were susceptible to ferroptosis we treated the cultures with iron and RSL3 (iron + RSL3), an inhibitor of GPX4 and known inducer of lipid peroxidation and ferroptosis [13]. While iron alone led to minimal cell death as assessed by Draq7 integration, a dye that permeates dead cells, inhibition of GPX4 induced robust cell death 20 hours post-treatment, suggesting that this culture system can undergo ferroptosis. Interestingly, RSL3 alone did not induce cell death suggesting that this human model system requires iron supplementation (Fig. S1B); these data are different than studies in immortalized cell lines in which RSL3 alone was sufficient to cause death [13, 20]. In support of a ferroptotic mechanism, the iron + RSL3 induced cell death was inhibited to iron alone (42.9% ± 14.8% SE of iron + RSL3) levels by two commercial ferroptosis inhibitors Ferrop_Inh1_ (63.6% ± 20.6% SE of iron + RSL3) (p<0.05) and Ferrop_Inh2_ (40.9% ± 11.6% SE of Iron + RSL3 (P<0.01) (Fig. 1D). These results suggest that we can utilize this human tri-culture system to study the role of ferroptosis.

**Figure 1:**
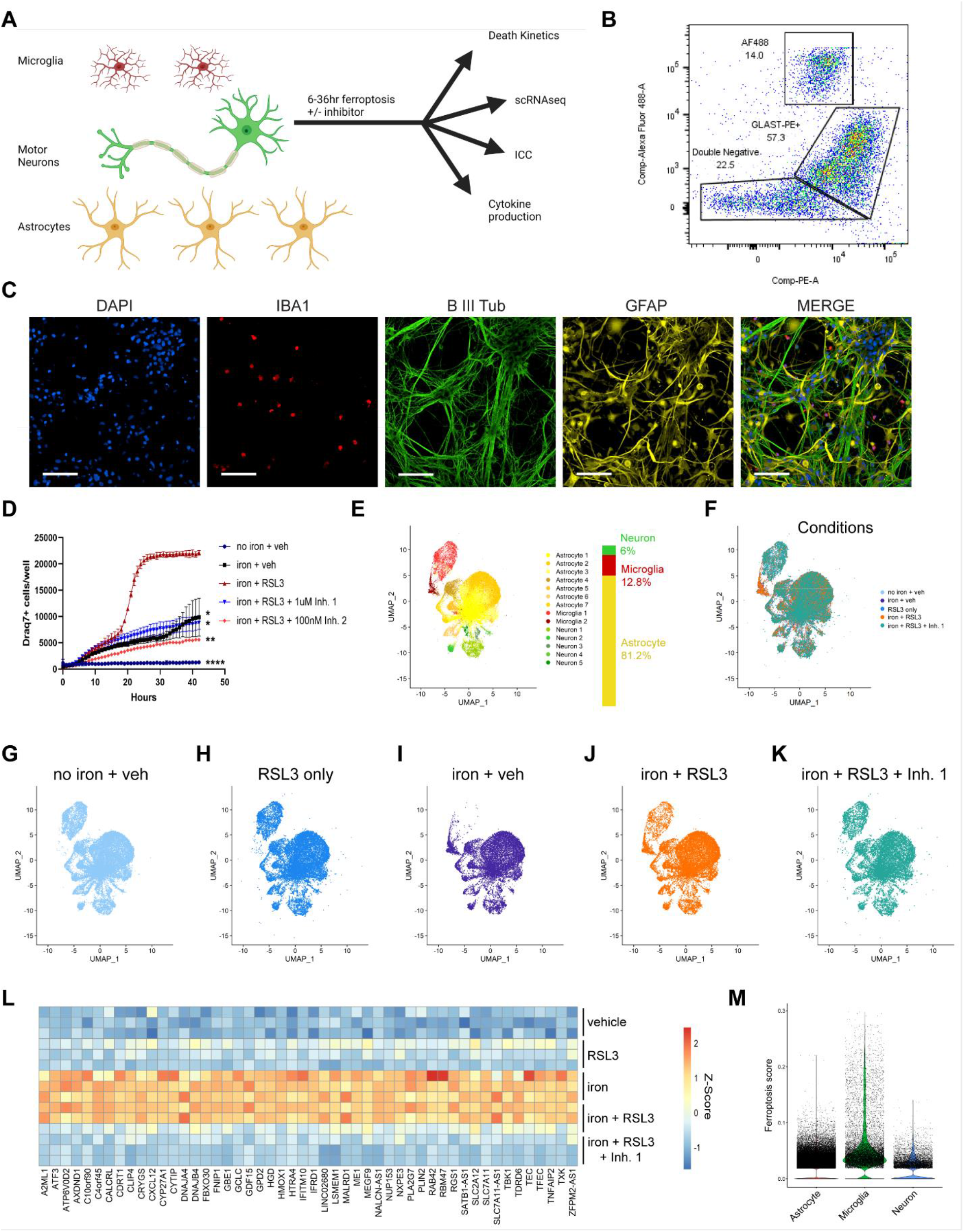
Ferroptosis induction causes a unique transcriptional response and cell death in iPSC tri-cultures. **(A)** Schema for generation of tri-culture and downstream analysis. **(B)** Flow cytometry analysis isolating GFP+ microglia, GLAST-PE+ astrocytes, and double-negative neurons. (n=1). (**C**) Representative image of tri-culture showing IBA1+ (red) microglia, βIII-tubulin+ (green) neurons, and GFAP+ (yellow) astrocytes. Scale bar = 100μm. (**D**) Draq7+ death kinetics in tri-cultures exposed to 1600uM iron + 1uM RSL3 ± commercial ferroptosis inhibitors. (n=4). Representative graph. AUC, log transformed. two-way ANOVA, Dunnett post hoc. *p<0.05, **p<0.01, ****p<0.0001. Error bars represent SEM. **(E)** UMAP representation of single cell RNA seq analysis of 108,456 cells from tri-cultures exposed to 1600uM iron + 1uM RSL3 ± commercial ferroptosis inhibitors. (**F-K**) UMAP colored by treatment condition **(L)** Pseudobulk analysis of all cell types and heatmap of top dysregulated genes. (n=3). **(M)** Violin plot of gene signature enrichment UCell score for all three cell types using genes from **(L)**.

To elucidate how each cell type responded to ferroptosis induction, we investigated cell-specific transcriptomic changes in the tri-culture system by performing scRNAseq. We analyzed the cells 6 hours after ferroptotic stimulation, a timepoint that precedes significant death induction in the iron + RSL3 condition (Fig. 1D). Transcription inhibitors were used to preserve cell state during the preparation of the cells for sequencing [21]. Altogether, 108,455 cells were sequenced with an average read depth of ~23,000 counts per cell. We identified 13 clusters across the five conditions: Vehicle treated (no iron + veh), RSL3 only, iron only, iron + RSL3, and iron + RSL3 + Ferrop_Inh1_ (Fig. S1C). Using cluster gene expression signatures and several known markers for each cell type, we were able to clearly distinguish the microglia (12.8%), neurons (6%), and astrocytes (81.2% (Fig. 1E, S1D-F). We performed unbiased pseudo-bulk analysis and identified genes that were induced in the iron + RSL3 condition that were also reversed by Ferrop_Inh1_ (Fig 1L). This unbiased ferroptosis-associated signature (FAS) was then applied to examine enrichment through UCell scoring [22] for each cell type. We found that microglia had the strongest induction of the ferroptosis signature, which was supported by the presence of a unique FAS microglia sub-population in the iron only and iron + RSL3 conditions (Microglia 2, Fig. 1E–K). This FAS microglial-specific cluster was almost nonexistent in the no iron + veh control and RSL3 only conditions, and it was markedly reduced in the iron + RSL3 + Ferrop_Inh1_ condition (Fig. 1G, H, K, and S1G). Our data shows a robust alteration in the transcriptional profile of the microglia prior to ferroptosis suggesting that these cells maybe the most sensitive responders to iron (Fig. 1M, S2A-C).

### Neurons and Astrocytes have a more subtle ferroptosis induction

To further understand the ferroptosis-dependent signature, we examined differentially-expressed genes (DEGs) between the iron + RSL3 and no iron + veh conditions in each cell type (Fig. 2A–C). FAS microglia upregulated several ferroptosis-related genes including ferritin, *FTH1*, and the glutathione-related genes *SLC7A11* and *GCLM* (Fig. 2A), which are necessary to produce glutathione, the main reducing agent for lipid peroxidation in ferroptosis [23, 24]. KEGG pathway analysis [25] of the top 50 differentially expressed genes showed that ferroptosis was the most significantly affected pathway. Glutathione metabolism was also one of the top associated pathways as well as MAPK signaling, which has been implicated in ferroptosis induction through the voltage-dependent anion channel (VDAC) in the mitochondria [26–28] (Fig. 2D). Several of the same ferroptosis and iron-related genes that were upregulated in the FAS microglia were also differentially expressed in astrocytes and neurons, including *SLC7A11*, *GCLM*, and *TFRC* (Fig. 2B and C). Despite this similarity, the number of dysregulated genes was much lower in neurons and astrocytes. Unlike microglia, the ferritin genes *FTH1* and *FTL* were both downregulated in neurons (Fig. 2B) suggesting cell-type differences in iron sequestration in the early stages of ferroptosis induction. However, ferroptosis was still one of the top affected pathways in neuron and astrocytes (Fig. 2E and F). These results indicate that neurons and astrocytes may have a delayed or muted response to ferroptosis-inducing stimuli compared to microglia.

**Figure 2:**
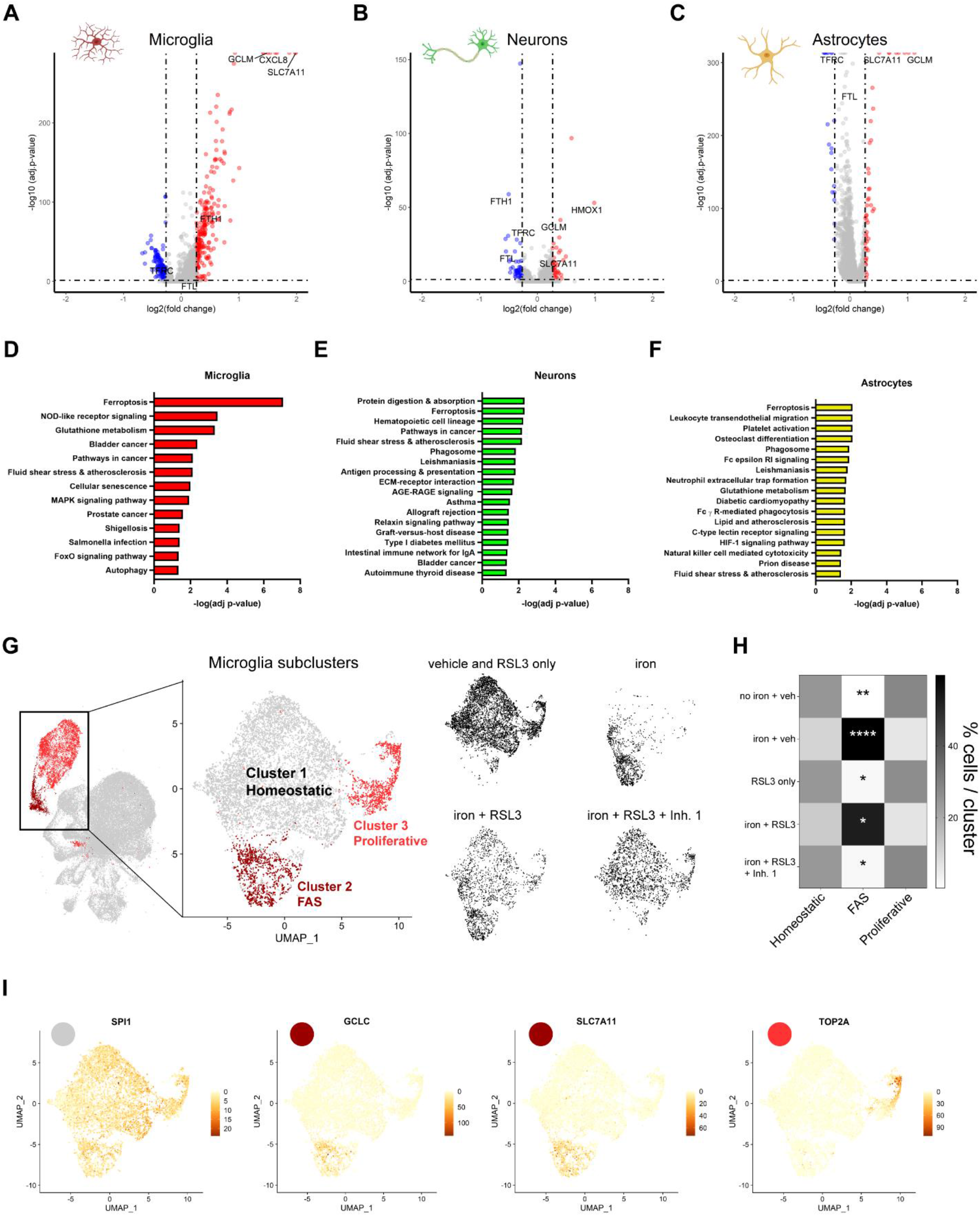
Ferroptosis induction causes a profound shift in microglia cell state compared to astrocytes and neurons. **(A)** to **(C)** Volcano plot of transcriptional changes in **(A)** microglia (**B**) neurons and (**C**) astrocytes in the iron + RSL3 condition versus no iron + veh. **(D)** to **(F)** KEGG pathway analysis of top upregulated and downregulated genes in the **(D)** microglia, **(E)** neurons, and **(F)** astrocytes in the iron + RSL3 condition versus no iron + veh. **(G)** UMAP microglia subclusters (homeostatic, ferroptosis-associated signature (FAS), and proliferative) and plots for each treatment condition in black. **(H)** Quantification of microglia subclusters per condition. Two-way ANOVA, Tukey post hoc. *p<0.05, **p<0.01, ****p<0.0001. **(I)** UMAP gene expression plots for the microglia marker *SPI1* (grey in **(G)**), the ferroptosis markers *GCLC* and *SLC7A11* (maroon in **(G)**), and the cell proliferation marker *TOP2A* (pink in **(G)**).

To understand the effect of the ferroptotic stimuli directly on microglia cell-state we subclustered the microglia and uncovered 3 unique subpopulations (Fig. 2G). We identified cluster 2 as the iron-induced FAS microglial cluster; the cells in this cluster expressed the iron and ferroptosis-related genes *GCLC* and *SLC7A11*. We also identified cluster 1 as the homeostatic population most represented in the control conditions and cluster 3 as a proliferative subset of microglia by expression of *TOP2A* (Fig. 2G–I). The FAS microglia cluster was significantly enriched in the iron (p<0.0001) and iron + RSL3 (p<0.05) conditions and significantly reduced in vehicle (p<0.01), RSL3 only (p<0.05), and iron + RSL3 + Ferrop_Inh1_ (p<0.05) conditions (Fig. 2G and H). There were no changes in the proportion of microglia in the proliferative subpopulation. These data demonstrate that microglia undergo a drastic shift in cell state following exposure to iron and prior to cell death, suggesting a functional consequence of iron-overload in these cells.

### Microglia uptake iron and secrete the inflammatory cytokine IL-8

Our data suggest that transcriptionally, microglia are most affected by alterations in iron homeostasis but does not address whether this also manifests in a functional alteration. One of the top differentially expressed genes in the FAS microglia was the gene encoding the ferritin heavy chain, *FTH1*. Ferritin is the main protein that sequesters iron in the cell and increased expression has been associated with ferroptosis in a disease context [29]. *FTH1* was significantly upregulated in the FAS microglia (Fig. 2A and 3A). We performed immunocytochemistry to confirm increased expression, as well as compare protein expression across cell types. Consistent with the transcriptomic data, FAS microglia increased expression of ferritin by 2.3-fold over vehicle treated cells (p<0.05). Astrocytes also increased expression over vehicle treated, but to an overall lesser magnitude, with iron + RSL3 treated microglia producing 2.9-fold more ferritin than iron + RSL3 treated astrocytes (Fig. 3C and D). This finding corroborates that microglia play a major role in sequestering iron [9, 30].

**Figure 3:**
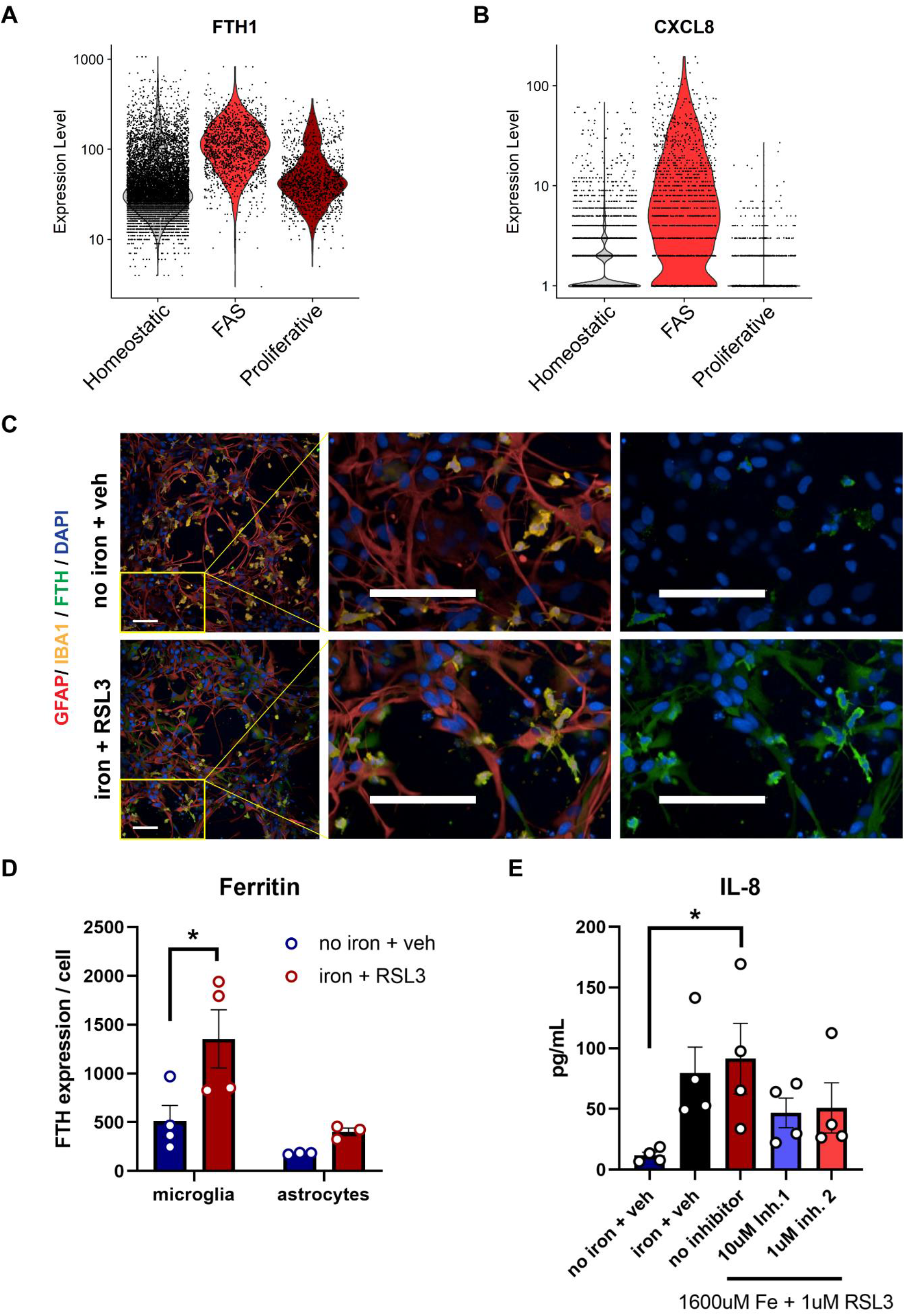
Microglia produce the majority of ferritin and increase IL-8 production during ferroptosis. **(A)** and **(B)** Violin plots for *FTH1* and *CXCL8* in the homeostatic microglia, ferroptotic microglia, and proliferative microglia. (**C**) Representative images of FTH1 (green) expression in IBA1+ microglia (yellow) and GFAP+ astrocytes (red) in tri-cultures 18 hours post-treatment. Scale bar = 100μm. (**D**) Average expression of FTH1 per IBA1+ microglia (n=4) or GFAP+ astrocyte (n=3). Log transformed, two-way ANOVA, Sidak post hoc. *p<0.05. Error bars represent SEM. **(E)** IL-8 production among conditions (n=4). One-Way ANOVA, Dunnett post hoc. *p<0.05. Error bars represent SEM.

These data suggest that microglia are highly sensitive to iron homeostasis, but to address whether this also leads to altered downstream signaling, we examined whether microglia change their secretory profile following iron challenge. We measured cytokine production from the supernatants in the human tri-culture system using a multiplexed approach. We identified a marked 7.8-fold increase in secreted IL-8 (p<0.05) (Fig. 3E) while the other 9 cytokines tested were not detectable, suggesting a targeted inflammatory response. Interestingly, *CXCL8* the gene encoding IL-8, was one of the top upregulated genes in the FAS microglia with a 2.6-fold increase (Fig. 2A and 3B). IL-8, which serves as a chemoattractant, has been linked to neurodegenerative disorders including ALS and PD [31]. There was a stepwise increase in IL-8 levels from iron only to Iron+RSL3. Remarkably, treatment with Ferrop_Inh1_ and Ferrop_Inh2_ partially blocked increases in IL-8 production (Fig. 3E). These results suggest the microglia are producing a specific inflammatory response that can be reduced by blocking ferroptosis. Overall, these results suggest that microglia are the major cell type to uptake iron and the first to induce ferroptosis (Fig. S2A-C) and produce an inflammatory response that could contribute to the pathological environment in disease.

### Microglia exhibit lipid peroxide ferroptosis signature

Our data thus far show that microglia are sensitive to iron handling by undergoing phenotypic changes and cell death. Furthermore, we show that commercial ferroptosis inhibitors can block both the signaling alterations and cell death. However previous work has shown that ferroptosis is induced by a specific set of arachidonic acid-derived lipid peroxides that leads to loss of membrane integrity and cell death.[20, 32]. Therefore, we sought to determine if human microglia treated with iron + RSL3 were susceptible to lipid peroxidation. We utilized an immortalized microglia cell line derived from primary, human adult microglia which is susceptible to ferroptosis and blocked by commercial inhibitors [11] (Fig. S3A). We performed lipidomic analysis at 2 hours post induction before cell death (Fig. S3B) and measured free 12-HETE and 15-HETE, which are the reduced forms of the hydroperoxides 12-HpETE and 15-HpETE. Indeed, ferroptosis induction caused a 2-fold increase in 12-HETE (p<0.0001) and a 30-fold increase in 15-HETE (p<0.0001), which was prevented with Ferrop_Inh2_ co-treatment. Interestingly, neither iron alone nor RSL3 alone was able to significantly induce these lipid peroxides (Fig. S3C and D). This is consistent with the data obtained in the tri-culture system (Fig. 1D and S1B) that both iron and RSL3 are necessary to fully induce ferroptosis. Iron alone seems sufficient to induce the ferroptotic gene signature (Fig. 1L), but inhibition of GPX4 is necessary to fully induce ferroptosis. A specific oxidized lipid, 1-SA-2-15-HpETE-PE, has been implicated in ferroptosis and was increased in models of PD and in fibroblasts from PD patients [32]. In the microglia we uncovered a 6-fold increase in 1-SA-2-15-HpETE-PE (p<0.01), which was also blocked with Ferrop_Inh1_ or Ferrop_Inh2_ treatment (Fig. S3E). These findings show that the microglia produce the defined, distinct ferroptotic lipid hydroperoxyl signature.

### FAS microglia are present in human neurodegenerative disease

There is strong evidence for the involvement of ferroptosis in many neurodegenerative diseases, including ALS and PD [7, 32–35]. However, identifying a consistent signature has remained elusive, and there has been little focus on immune cells and microglia. Inflammation from microglia and peripheral immune cells is known to play a significant role in disease progression. We wanted to determine if we could identify a shared ferroptosis transcriptomic signature across multiple neurodegenerative diseases. Iron dysregulation has been identified in ALS patients, including increased iron accumulation in the deep cortical layers, as well as in microglia in the motor cortex, and iron chelators can increase life expectancy in a mutant SOD1 mouse model of ALS [5, 7]. Iron overload can lead to oxidative stress and death through pathways independent of ferroptosis [36]. Thus, we wanted to determine if the ferroptotic signature identified in the tri-culture FAS microglia was present in ALS patient tissue. We compared gene expression changes between ALS patient and case controls from the Target ALS consortia (Table S1) for the top 50 upregulated genes in the FAS microglia. Indeed, we found significant upregulation of the gene set especially in the spinal cord of patients, including *SLC7A11* and *GCLM*, which were upregulated across all cell types in the tri-culture. To a lesser degree, there was also dysregulation in the motor cortex, frontal cortex, occipital cortex, and cerebellum (Fig. S4A). These results demonstrate a strong ferroptotic signature in ALS patients, primarily in the spinal cord, suggesting ferroptosis induction in the region most affected by disease.

Previous work identified a subset of microglia in MS patients with an iron-related / ferroptotic signature [11, 12, 37]. To determine if the iron-related signature found in MS microglia is present in PD, we analyzed snRNAseq of putamen tissue from three PD patients and three healthy controls (Fig. 4A) (Table S2). The putamen is a pathologically relevant area in the mid brain for PD, with significant connections to the substantia nigra and reduced spontaneous activity leading to impaired task performance in PD patients [38, 39]. Unbiased clustering identified microglia, oligodendrocyte precursor cells (OPCs), oligodendrocytes, neurons, astrocytes, and endothelial cells in the sequenced samples (Fig. 4B and S5A). To determine which cell type might exhibit a ferroptotic signature we performed differential expression analysis of the MS microglia iron and ferroptosis-related gene signature across all cell types and found that the microglia in PD tissue were uniquely enriched for the gene signature (Fig. S5B). To further investigate the signature, we identified six microglia subpopulations in the dataset (Fig. 4E and S5C). Microglia subcluster 1 had the most prominent MS ferroptosis signature (Fig. S5D). This gives further evidence, at the transcriptomic level, that ferroptosis may be occurring in the microglia of PD patients.

**Figure 4:**
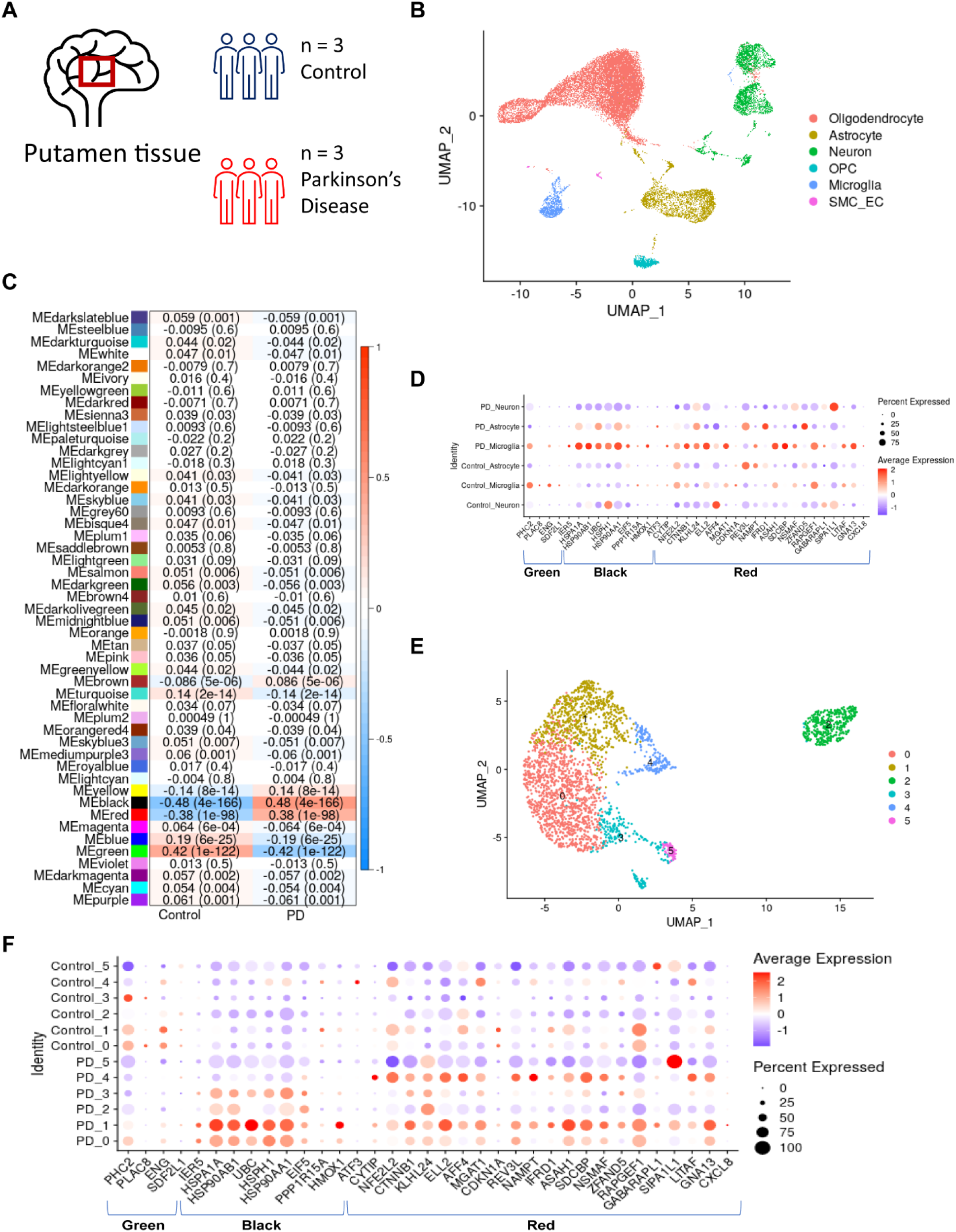
Ferroptosis signature identified in the microglia of PD patients by snRNAseq. **(A)** Brain region and sample size for single Nucseq dataset. **(B)** Unsupervised clustering and annotation of cell types identifies microglia cluster present in control and PD samples. **(C)** WGCNA analysis uncovers three modules significantly associated with the PD microglia (green, red, and black). (**D**) Dot plot of shared genes in green, red, and black modules with microglia tri-culture signature in different control and PD cell types. **(E)** Subclustering of microglia from **(B)** to identify subpopulations. **(F)** Dot plot of shared genes in green, red, and black modules in microglia subpopulations.

To further investigate the ferroptotic signature in PD microglia, we utilized our tri-culture ferroptotic microglia signature, which we found to be enriched in ALS spinal cord (fig. S3A). We performed weighted gene co-expression network analysis (WGCNA) on the control and PD snRNAseq dataset (Fig. S5A and B). This analysis revealed the 3 modules that were most significantly correlated with PD: red [R=0.38, p=1×10^−98^], green [R=-0.42, p=1×10^−122^], and black [R=0.48, p=4×10^−166^] (Fig. 4C, S6A and B). The red and black modules were positively correlated with PD and were enriched for our tri-culture-derived ferroptosis signature genes including *HMOX1* and *CXCL8* suggesting that PD patients show enrichment of a ferroptosis signature. Interestingly, when we compared the genes from those modules across neurons, microglia, and astrocytes from the snRNAseq datasets, we found the highest level of differential expression in the microglia (Fig. 4D). The red and black module genes were more enriched and highly expressed in PD microglia than in control microglia. Genes from the green module, which was anticorrelated with PD, showed increased gene expression in the control patient microglia relative to PD patient microglia and could demonstrate dampening of a homeostatic microglial signature in PD patients. We then analyzed the microglia subclusters (Fig. 4E) for the ferroptosis-associated signature and, like the MS microglia signature, PD microglia subcluster 1 had the strongest upregulation of the red and black module genes. However, the red and black modules were also enriched in subclusters 0, 2, 3, and 4. These results show a strong, shared ferroptosis-associated signature in microglia specifically in disease-afflicted areas across multiple neurodegenerative disorders including MS, PD, and ALS.

### Analysis of PD patient blood reveals ferroptotic signature

Our data thus far indicate that iron dysregulation leads to a distinct signature in microglia. We next sought to investigate whether there might be evidence of altered iron homeostasis systemically in patients with neurodegenerative disease. For this, bulk RNA-Seq analysis was performed on blood samples obtained from case and control participant enrolled in cohort studies of the Accelerating Medicines Partnership Parkinson’s Disease (AMP PD), the Parkinson’s Progression Markers Initiative (PPMI) (n=1,433; 816 PD case: 617 control), and Parkinson’s Disease Biomarkers Program (PDBP) (n=1,284; 780 PD case: 504 control) (Fig. 5A) (Table S3). Unbiased differential gene expression analysis between PD cases and healthy controls uncovered significantly up- and down-regulated ferroptosis-related genes, many of which were found in our tri-culture FAS microglia signature including, *GCLC*, *GPX4*, *HMOX1*, *SQSTM1*, and *SLC7A11* (Fig. 5B) (Table S4). Indeed, ferroptosis and IL-8 signaling were among the top dysregulated pathways identified using Ingenuity Pathway Analysis (IPA) (Fig. 5C). These findings suggest that a ferroptotic gene signature is present in PD patient-derived blood samples and that iron dysregulation may be present systemically in patients. Furthermore, these findings show that iron dyshomeostasis is observed peripherally in PD and that blood samples could potentially be used as a peripheral biomarker for central nervous system (CNS) ferroptosis pathway activation in PD and other neurodegenerative diseases.

**Figure 5:**
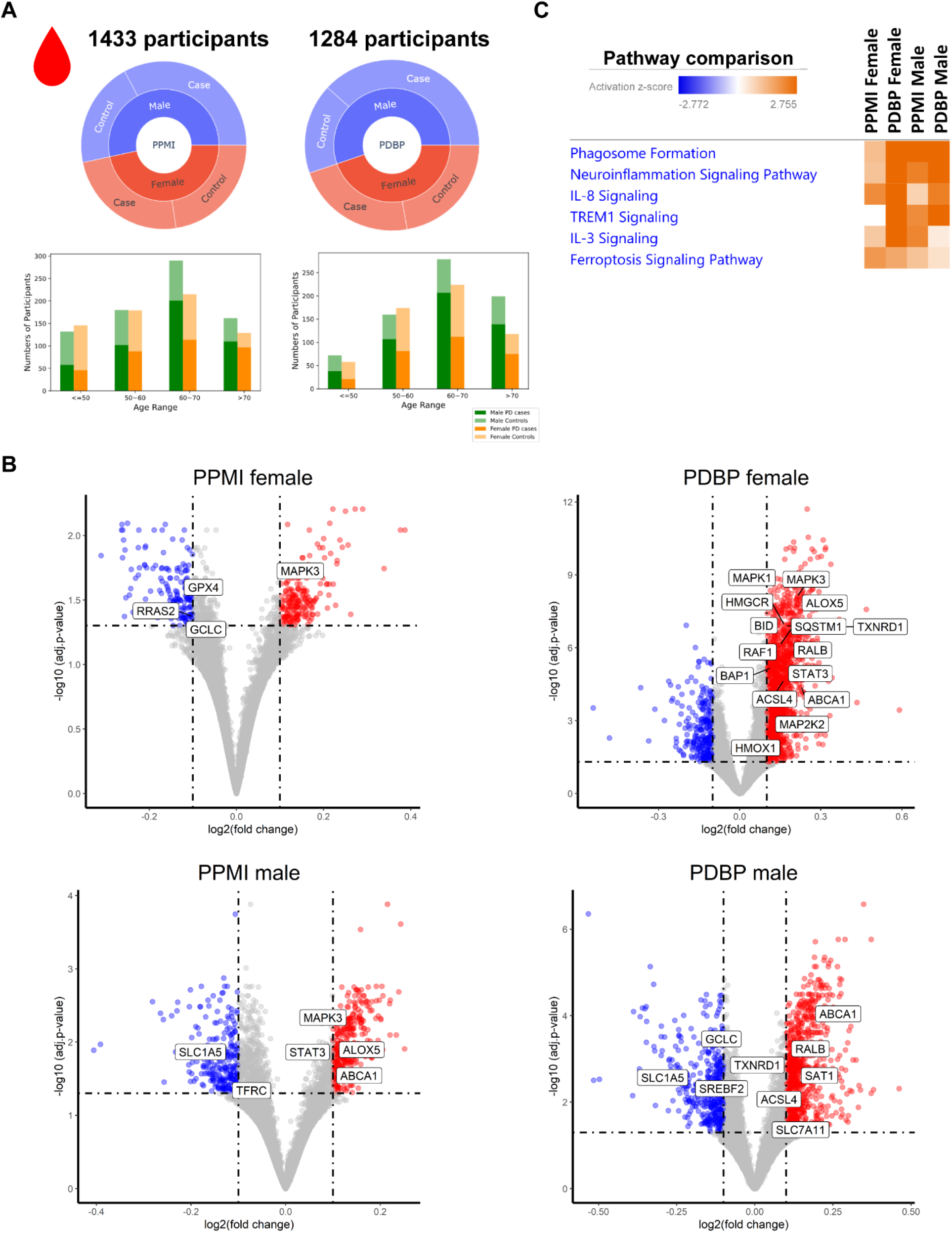
Detection of a ferroptosis gene signature in the AMP-PD Parkinson’s patient blood samples. **(A)** Breakdown of case, control, age, and gender for the PPMI and PDBP studies by the AMP-PD consortia. **(B)** Volcano plots for DEGs, −0.1<log2 fold change < 0.1 and adjusted p-value < 0.5, between healthy controls and PD patients stratified by gender and cohort. **(C)** IPA Comparison Analysis of differentially expressed genes (inclusion cutoffs: −0.1<log2 fold change < 0.1 and adjusted p-value < 0.5 with male and female patients in PPMI and PDBP studies).

### SEC24B regulates ferroptosis in microglia

Susceptibility to ferroptosis varies among cell types, [40–42] so we sought to determine if there are unique regulators of ferroptosis in microglia. Genome-wide screens for ferroptosis regulators have been performed in several cancer cell lines and have identified regulators including *FSP1*, *POR*, and *ACSL4* [43–47]. However, little is known about ferroptosis regulation in brain cells. We performed a positive selection genome-wide CRISPR screen using Cas9-expressing immortalized human microglia. We utilized a 76,612-guide library that targets 19,114 genes and uses 4 guides per gene. Cells were transduced with two separately generated viral pools and treated with vehicle or iron + RSL3 for 8 or 24 hours to have shorter or longer selection, respectively (Fig. 6A). We found 61 hits from the 8 hour exposure (Fig. 6B). Hits were stratified by the number of guide (g)RNAs that were detected for each gene in the surviving pool with p values < 0.05 (Fig. 6C and S8A and C). Pathway analysis using PANTHER [48] identified several necrosis and necroptotic pathways as the most significantly associated with the 8 hour hits (Fig. S8B). Furthermore, 65 hits were found in the 24 hour condition (Fig. 6B). Interestingly, these hits were associated with different pathways than those found in the 8hr condition, with G-protein coupled receptor signaling being the most significantly associated pathway (Fig. S8D).

**Figure 6:**
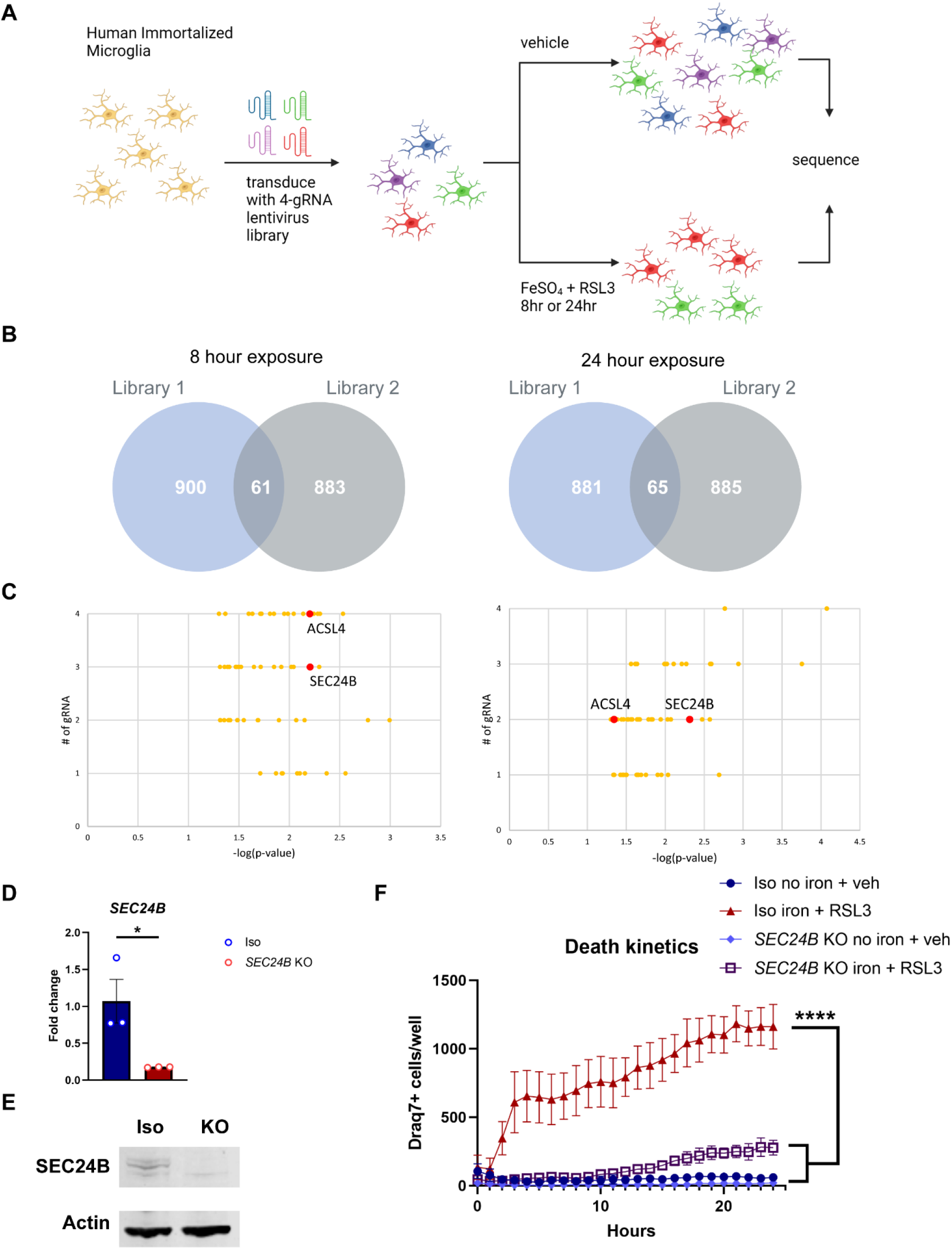
Genome-Wide CRISPR screen identifies *SEC24B* as a novel regulator of ferroptosis in microglia. **(A)** Schema for positive selection Genome-wide CRISPR screen in Cas9-expressing human immortalized microglia. **(B)** Venn diagrams for hits from each viral pool in 8hr and 24hr treatments. **(C)** Overlapping hits from each viral pool with number of gRNAs identified in 8hr and 24hr treatment with *SEC24B* and *ACSL4* highlighted. **(D)** qRT-PCR analysis showing markedly reduce expression of *SEC24B* in KO line (n=3). Unpaired t test. *p<0.05. Error bars represent SEM. **(E)** Western blot showing absence of SEC24B protein in KO line. **(F)** Death kinetics in *SEC24B* KO Hap1 cell line and isogenic control (n=3). AUC, one-way ANOVA, Dunnett post hoc. ****p<0.0001. Error bars represent SEM.

Across the 126 hits, *ACSL4* and *SEC24B* were present in both timepoints and libraries (Fig. 6C). *ACSL4* has been identified in several previous genome-wide CRISPR screens [43–45], but *SEC24B* has not been previously described as a regulator of ferroptosis. SEC24B is a COPII coat complex component that is important for vesicle trafficking from the endoplasmic reticulum (ER) to the Golgi apparatus [49]. To confirm SEC24B as a regulator of ferroptosis we assessed ferroptosis susceptibility in an *SEC24B* KO myeloid cell line. SEC24B knockout was confirmed by qRT-PCR, which showed a roughly 80% reduction in RNA expression (p<0.05), and an absence of protein by Western blot (Fig. 6D and E). Cells were incubated with Draq7 and ferroptosis was measured over 24 hours. The SEC24B cells were highly resistant to ferroptosis with a 4-fold reduction in ferroptosis as compared to the control, isogenic line (p<0.0001) (Fig. 6F). These results demonstrate that *SEC24B* is required for ferroptosis induction in microglia and macrophages.

### Pharmacological screen identifies clinical compounds that inhibit ferroptosis in microglia and corroborates CRISPR-identified pathways

To test whether we could also inhibit ferroptosis pharmacologically we performed a targeted small molecule screen and identified molecules that are already in clinical trials or can be used as tool compounds for further drug development. We utilized a commercially available, ferroptosis-related library that consists of 546 compounds, including some that are already in clinical use [50, 51]. Ferroptosis was induced in the human microglia cell line and were co-treated with the compounds at 10 μM or 2 μM. Inhibition of ferroptosis was noted for any compound that led to ≥70% cell viability compared to vehicle control. Of the 546 compounds we found 39 compounds that inhibited ferroptosis (Fig. 7A and B) (Table S5). Interestingly, 3 of the compounds, leveocetirizine dihydrochloride, jatrorrhizine, and 8-OH-DPAT, targeted neuronal signaling, and 2 of the compounds, olivetol and xanthotoxol, targeted G protein-coupled receptor (GPCR) signaling. Xanthotoxol targets the GPCR 5-HT receptor (Fig. 7C), which is involved in mediating excitatory and inhibitory neurotransmission [52]. The 5-HT receptor is also expressed on microglia and may be involved in inflammation [53].

**Figure 7:**
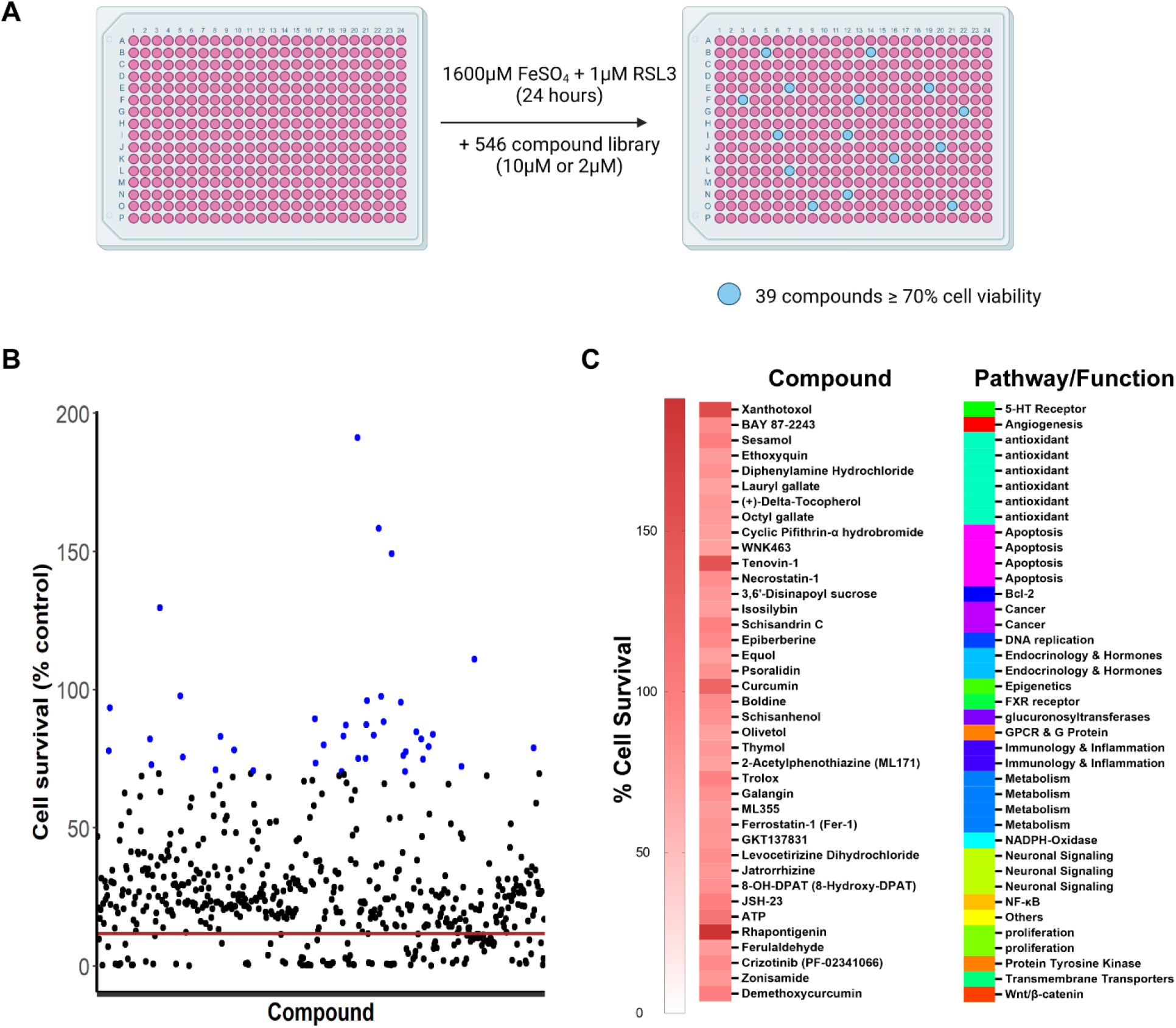
Ferroptosis pharmacological inhibitors revealed in small molecule screen. (**A**) Schema and results for pharmacological compound screen in which 546 compounds were tested for their ability to block ferroptosis in an immortalized microglia cell line. (**B**) 39 of the 546 compounds rescued cell viability to at least 70% of the vehicle control (blue dots) and 5 rescued to at least 100%. 3 technical replicates per compound. Red line indicates average cell survival with ferroptosis induction and no compound. (**C**) Heatmap of percent cell survival for the 39 hits and the associated pathway or function.

In addition to validating multiple pathways from the CRISPR screen, several of the other hits corroborated previously identified regulators of ferroptosis. Galangin targets cytochrome P450 and ML171 and GKT137831 both target NADPH-oxidase (Fig. 7C). Cytochrome P450 oxidoreductase, which utilizes NADPH, has been recently identified in a separate genome-wide screen [45]. Additionally, the p53 inhibitors cyclic pifithrin-α hydrobromide and Tenovin-1, were also effective at preventing ferroptosis. P53 has recently been identified as an alternative pathway for ferroptosis induction [42]. Lastly, BAY 87-2243 is a HIF-1α inhibitor in phase 1 clinical trials for cancer. HIF-1α was upregulated in the ferroptotic microglia in the tri-culture. These results reaffirm the pathways identified in the CRISPR screens and point to possible clinical compounds that could be used for the treatment of ferroptosis-related neurodegenerative diseases.

## DISCUSSION

While ferroptosis has been previously described in microglia [17], the relationship of ferroptosis induction to neurodegeneration has not been explored. Iron overload in microglia has been well described [7, 11, 54] but the consequences of this overload at the transcriptomic and functional level are not well understood. Here, we identify microglia as a major player in a ferroptotic cascade during neurodegeneration. In the tri-culture model, the microglia are the first to develop a ferroptotic signature. Microglia may act as an initiator of ferroptosis and/or inflammation via IL-8 production that leads to neurotoxicity [31]. Additional temporal studies would be required to confirm the microglia-mediated neuronal death in response to iron rather than cell autonomous neuronal ferroptosis. It is possible that microglia play a protective role by sequestering iron, but as they undergo ferroptosis, iron is released into the extracellular space and taken up by other cell types. In support of this mechanism, in our tri-culture system, the neurons die after *en masse* death of the microglia.

We found that the ferroptosis-associated signature (FAS) correlates well across multiple neurodegenerative diseases. We and others have previously identified a ferroptotic signature in MS microglia [11, 12]. Using snRNA-Seq, we established a unique ferroptotic gene signature in our tri-culture system microglia. This ferroptotic gene signature was enriched in the ALS patient spinal cord, as well as in microglia from PD patient putamen. This supports the utility of our human iPSC-derived tri-culture system to recapitulate disease-relevant ferroptosis signatures and the activation of ferroptosis pathways in ALS and PD specifically in microglia in disease-associated tissues. Determining when and where ferroptosis occurs and identifying biomarkers that could help stratify patients will be beneficial for treatment in the clinic. Existing efforts to identify individual markers of ferroptosis has been difficult [55], and here we demonstrate that a transcriptomic signature could be another useful tool. We identified a ferroptotic signature in two separate tissues, blood and brain, suggesting that there could be several approaches used to identify affected patients.

Regulation of ferroptosis in microglia has not yet been studied. To this point, no genetic or pharmacological screen for ferroptosis regulators has been performed on microglia. Using a genome-wide CRISPR screen, we discovered that, in addition to the well-described regulator *ACSL4*, *SEC24B* also strongly regulates ferroptosis in microglia. SEC24B has never previously been implicated as a regulator of ferroptosis. Under homeostatic conditions, it is involved in vesicle trafficking from the endoplasmic reticulum (ER) to the Golgi apparatus, particularly for secretory proteins [56]. ER stress has been implicated in ferroptotic induction [26, 57]. This stress may prevent trafficking of anti-ferroptotic proteins such as GPX4, as reduced GPX4 expression has been identified in multiple neurodegenerative disorders including MS and ALS [58, 59]. Additionally, chronic iron overload has been associated with impaired protein secretion [60]. Loss of *SEC24B* could dysregulate secretory proteins like transferrin, which is necessary for cellular uptake of iron. *SEC24B* could be a unique regulator of ferroptosis in microglia or at least in a subset of cell types. Future studies exploring expression and subcellular localization of ferroptosis-related proteins such as transferrin and ferritin as well as iron uptake and trafficking may help determine how SEC24B regulates ferroptosis.

Despite the fact that ferroptosis has been implicated in many disorders, an effective therapeutic to mitigate ferroptosis has not been developed for patients [36]. Iron chelators are one potential approach, but many have unknown or poor blood brain barrier permeability and may disrupt homeostatic redox functions [61]. Given the clear role of ferroptosis in neurodegenerative diseases, there is a need for a more exploration into ferroptosis-related therapeutics. There are several more iron-related compounds in clinical trials, including the vitamin E derivative vatiquinone, deuterated linoleic acid, and activators of the antioxidant NRF2 pathway. We screened a commercially available library of ferroptosis-related compounds in the human microglia cell line. We found several compounds involved in established ferroptotic regulatory pathways. This included the HIF-1α inhibitor BAY 87-2243, which is in phase 1 clinical trials for cancer, as well as the FDA-approved antioxidant Octyl gallate. These compounds could also be tested for efficacy in ferroptosis-related neurodegenerative disorders.

Overall, our work further confirms the role of ferroptosis in multiple neurodegenerative disorders, including ALS and PD. Using a unique tri-culture system, we show cell type-specific disease signatures of ferroptosis identifying microglia as an initiating cell type in ferroptosis. Through a genome-wide CRISPR screen, we also describe a novel regulator of ferroptosis, *SEC24B*, in microglia. Further understanding the role of SEC24B in microglia and ferroptosis may lead to new insights for therapeutic targets for treating multiple neurodegenerative disorders. Finally, we demonstrated pathway convergence of the genetic and pharmacologic screens on pathways that drive ferroptosis, furthering the understanding of the molecular mechanisms of microglial ferroptosis. Altogether, our work supports the importance of iron dyshomeostasis in microglia as a critical driver across multiple neurodegenerative diseases.

## MATERIALS AND METHODS

### Study Design

In this study, to understand the role of ferroptosis in neurodegenerative disease, we aimed to evaluate the susceptibility of disease relevant cell types to ferroptosis. We developed a tri-culture of hiPSC-derived neurons, astrocytes, and microglia and placed cultures under ferroptotic conditions (iron + RSL3) ± ferroptosis inhibitors. To understand the susceptibility to ferroptosis of each cell type, gene expression changes were measured by single cell RNAseq, protein expression changes were measured by ICC, and functional changes were measured by cytokine expression and death kinetics via Draq7 incorporation. These analyses identified microglia as highly susceptible to ferroptosis. Next, we investigated the microglia ferroptotic signature in multiple neurodegenerative disease patient samples through bulk RNAseq and single Nucseq, identifying the ferroptotic signature. We utilized a genome-wide CRISPR screen to identify regulators of ferroptosis in microglia and identified *SEC24B* and validated in a separate *SEC24B* KO myeloid cell line. Commercial compounds were screened for ferroptosis inhibition in a human microglia cell line to identify translatable chemical material.

### Study size calculations

Power calculations were not completed for these studies.

### Treatment of outliers

No outliers were removed for these studies.

### Randomization

Plate wells were randomly assigned to treatment groups.

### Blinding

Studies were not blinded. ICC and Draq7 quantification were automatically counted by the appropriate software.

### Replication

The number of replicates for each experiment and the test used to calculate statistical significance is indicated in the figure legends and/or methods. All cell culture experiments had one to three technical replicates per biological replicate. Technical replicates were averaged together per biological replicate.

### Tri-culture assembly

On D0, 96 well plates (Perkin Elmer 6055302 or Corning 3595) were coated with Matrigel (Corning 354277) (diluted in appropriate amount of DMEM (Life Technologies 11330057) solution). 200uL sterile PBS (Thermo Fisher Scientific 20012-027) was added to any unused wells. iAstrocytes (Fujifilm ASC-100-020-001-PT) were thawed and plated at 1.5×10^4^ per well in Fuji designated astrocyte media (200uL per well) (DMEM/F12, HEPES (Life Technologies 11330057) + 2% heat inactivated fetal bovine serum Certified One Shot (Gibco A38400-01) + 1× N-2 supplement, 100x (Gibco 17502-048)). On D1, 3.5×10^4^ iCell motor neurons (Fujifilm C1048) were added per well. Motor neurons were thawed and resuspended at 3.5×10^4^ cells/200uL in complete Fuji Motor neuron media (100mL iCell Neural base medium 1 (Fujifilm M1010) + 2mL iCell Neural supplement A (Fujifilm M1032) + 1mL iCell Nervous system supplement (Fujifilm M1031)). Fuji astrocyte media was fully aspirated and 200uL of motor neuron cell suspension was added to each well. On D3, a 75% media exchange (150uL) was performed in all wells with NB/B27+ media (B-27 Plus Neuronal system kit (Life technologies A3653401). On D5, iCell Microglia (Fujifilm C1110) were pated at 1×10^4^ per well in NB/B27+ with iMg growth factors (B-27 Plus Neuronal system kit (Life technologies A3653401) + 25ng/mL M-CSF (peprotech 300-25) + 100ng/mL IL-34 (peprotech 200-34) + 50ng/mL TGF-β1 (peprotech 100-21). iMicroglia were thawed and resuspended in NB/B27+ with iMg growth factors at 1×10^4^ cells/100uL. 100uL of media was removed from each well and 100ul of iMicroglia cell suspension was added. Half media exchanges were performed on D7, D9, D11, and D14. All treatments were added on D15.

### Tri-culture ferroptosis treatments

2x solutions were made for all tri-culture treatments. Half media exchanges with 2x treatments were performed. Final concentrations were 1:1000 DMSO (Sigma D2650), 1600uM FeSO_4_ (Sigma F8633), 1uM RSL3 (Sigma SML2234), 10uM cayman ferroptosis inhibitor (Ferrop_Inh1_) (Cayman Chemical 10010468), and 1uM liproxstatin-1 (Ferrop_Inh2_) (Sigma SML1414). Cultures were treated for 6-42hrs depending on experiment. All inhibitors were added as co-treatments.

### scRNAseq cell preparation

6hrs post treatment, cells were washed 2x with PBS. Cells were then treated with 0.25% Trypsin + EDTA (Sigma T4049) + transcription/translation inhibitors (5ug/mL actinomycin-D (Sigma A1410) + 10uM triplotide (Sigma T3652) + 27.1ug/mL anisomycin (Sigma A9789) for 6min in 37°C 5% CO_2_ incubator. 1 volume of PBS + 2% FBS (Gibco A38400-01) + transcription/translation inhibitors + 1:100 DNAse was added to quench. Cells were combined from two wells and put through 40um cell strainers (BD Falcon 352235). Strained cell suspension was placed in a 1.7ml eppendorf tube on ice. Cells were counted by hemocytometer. Cell suspensions were spun at 4°C for 5min at 1500RPM. Supernatant was removed and cells were resuspended at 1,000 cells/uL in ice cold PBS.

### ScRNAseq

Cells were loaded onto the 10x Genomics Next GEM Single Cell 3’ V3.1 sample chip and run following manufacturers protocols. All 15 samples were processed in parallel and were amplified 11 cycles during the initial cDNA amplification and the 12 cycles for the sample index PCR. The samples were sequenced on an Illumina Novaseq 6000 to an average depth of approximately 40,000 reads/cell. Sequencing files were processed using the Cell Ranger 4 pipeline and GRCh38 human reference genome.

### ScRNaseq analysis

Samples were analyzed using Seurat 4.0.2 [62] in R version 4.0.3. Samples were filtered based on the overall distribution of UMIs and gene counts per cell to limit the inclusion of low-quality cells and doublets. Cells with less than 3500 genes and 8000 UMIs and greater than 9000 genes, 50000 UMIs, and 10% mitochondrial genes were removed from analysis. Following filtering, the variable features were selected using the “vst” method and using 2000 features. RunICA was performed for 75 ICs and then the FindNeighbors function was run using 33 ICs to build the KNN graph. Cluster resolution was set to “0.3” and the UMAP was generated using the RunUMAP function and using 33 ICs. Cell types were assigned by identifying genes unique to each cluster and through cross-referencing to known markers of each cell type and existing published datasets. UMAP plots and gene expression plots were generated using built in Seurat/ggplot2 plotting functions unless otherwise described. Cluster specific genes in figures 1 and 2 were generated using the FindAllMarkers function in Seurat. Ferroptosis-related genes for figure 3 were calculated using the FindMarkers function in Seurat and using DeSeq2 as the differential expression test comparing the vehicle and Iron+RSL3 conditions. Volcano plots were generated using ggplot2 and significant points were colored using log fold change > 1.2 and adjusted p value < 0.05 as cutoffs.

### Pseudobulk analysis

For the unbiased analysis of ferroptosis-related genes (Figure 1L) pseudobulk gene matrices were generated for each sample by calculating the row sums for each gene. The samples were then all combined into a single gene counts matrix and analyzed using Deseq2 with the treatment condition as the variable of interest [63]. Genes with count sums equaling 0 were removed from the analysis. Statistically significant genes (Adjusted P value < 0.05) in the Control vs.

Iron+RSL3 groups and Iron+RSL3 vs. Iron+RSL3+’468 samples were identified and cross-referenced to find genes that were upregulated in the stimulated group and downregulated in the inhibitor treated groups. Genes were plotted using the Pheatmap package in R.

### UCell analysis

To determine the cell type enrichment of the ferroptosis signature generated from the pseudobulk analysis the UCell package was used (Andreatta and Carmona, 2021). The UCell signature score is based on the Mann-Whitney U statistic and is agnostic to dataset cell type composition when assigning scores. The 57 overlapping genes from the pseudobulk analysis the gene signature input and used to calculate the UCell score for each cell in the dataset. UCell score violin plots were grouped by cell type (microglia, neuron, and astrocyte) and plotted using Seurat/ggplot2.

### Microglia subclustering analysis

Microglia were subset from the larger dataset based on cell type definitions presented in Figure 1 and reanalyzed using Seurat. The variable features were selected using the “vst” method and using 2000 features. RunICA was performed for 20 ICs and then the FindNeighbors function was run using 10 ICs to build the KNN graph. Cluster resolution was set to “0.05” and the UMAP was generated using the RunUMAP function and using 10 ICs. Microglia subtypes were identified by using the FindAllMarkers function in Seurat. The volcano plot of ferroptotic microglia markers was generated using ggplot2 and significant points were colored using log fold change > 1.2 and adjusted p value < 0.05 as cutoffs. UMAP plots and gene expression plots were generated using built in Seurat/ggplot2 plotting functions unless otherwise described.

To calculate the percent of cells per cluster from each sample, the number of cells from each sample in a given cluster was calculated and normalized to the number of cells per sample. These values were then normalized to the other replicate samples. Significantly enriched samples were identified using a two-way ANOVA with Tukey’s post hoc test and P values are reported in each figure.

### Cytokine analysis

Supernatants from tri-cultures were run on the Proinflammatory Panel 1 (human) (MSD K15049D) according to manufacturer’s instructions.

### Draq7 death kinetics in tri-culture

2x treatments described in the previous section had Draq7 (abcam ab109202) added at 1:150. Cells were treated as described and imaged once an hour on the incucyte S3. Images were analyzed on the incucyte software.

### Ferroptosis induction in HAP1 cell lines

*SEC24B* KO and isogenic control Hap1 cell lines (Horizon Discovery HZGHC001222c002 & C631) were plated at 2.5×10^4^ cells/ well of a 96 well plate in HAP1 media (Iscove’s Modified Dulbecco’s Medium (IMDM) (Gibco 12440-053) + 20% FBS (Gibco A38400-01). 24hrs post plating, cells were treated with 1600uM FeSO_4_ + 1μM RSL3 for 24 hours depending on the experiment. Cell death was tracked by Draq7 (abcam ab109202).

### Draq7 death kinetics for immortalized microglia and HAP1 cell lines

Draq7 (abcam ab109202) was added to treatments at 1:300. Cells were imaged in the incucyte S3 once an hour. Images were analyzed on incucyte software.

### Genome-wide CRISPR screen sgRNA lentiviral library production

Lenti-X 293T (Takarbio 632180) cells were thawed in complete Lenti-X 293T media (DMEM (Millipore D5796) + 10% Tet-Free FBS (Takarabio 631101) + 1x pen/strep (Millipore TMS-AB2-C)) on an uncoated T75 flask. Once cells reached 80% confluency, cells were re-plated at 1×10^6^ cells per 10cm dish in 8mL Lenti-X 293T media onto 10 dishes. Cells were grown to 80-90% confluence. Each dish was transduced with 1 vial of Guide-it Genome-Wide sgRNA Library Transfection Mix (Takarabio 632650) as per manufacturer’s instructions. 24hrs post-transduction, and additional 6mL of media was added to each dish for a total of 14mL per dish. 48hrs post-transduction supernatants were collected from each dish. Two dishes’ supernatants were pooled, creating 5 total libraries. 8mL of media was added back to each dish, and the supernatants were stored at 4°C. 72hrs post-transduction, the remaining 8mL of supernatant was collected from each dish and added to the respective stored supernatants for a total of 44mL per library. The libraries were centrifuged at 4°C at 500xg for 10 min. 200uL was removed to determine viral titer, and the remaining was aliquoted and stored at −80°C. Viral RNA was isolated with NucleoSpin RNA virus (Takarabio 740956) according to manufacturer’s instructions, provided in Guide-it CRISPR Genome-Wide sgRNA Library System (Takarabio 632646). Viral titer was determined with Lenti-X qRT-PCR titration kit (Takarbio 631235) according to manufacturer’s instructions.

### Genome-wide CRISPR screen

Human microglia cell line was plated in T150 flasks (BD Falcon 355001) at 20 million/flask in 25mL complete microglia media and placed in 37C 5% CO2 incubator. 24hrs post plating, puromycin (Takara bio 631305) was added at 5ug/mL. Cells were grown in 5ug/mL puromycin for 11 days. A full media exchange was performed without puromycin. 4 days post-removal of puromycin, Nunc non-treated T175 flasks (Thermo Scientific 159926) were coated with 9ug/mL retronectin (Takara bio T110B) and incubated overnight at 4°C. The following day, retronectin coating was removed, blocked in 2% BSA (Sigma A7030) for 30in at RT. Plates were washed 1x with PBS. Plates were coated with sgRNA pool at 60 MOI for 6hrs in 37°C 5% CO_2_ incubator. After 6hrs, viral supernatants were removed and cells were added at 1.67×10^7^ cells per flask across 6 flasks for a total of 1×10^8^ cells. Cells were placed in 37°C 5% CO_2_ incubator. 24hrs post-transduction, cells were treated with 100ug/mL hygromycin B (Takara bio 631309). Cells were exposed to 100ug/mL hygromycin B for 8 days. Cells were lifted as previously described and plated to 20 Nunc EasYFlask 75cm (Thermo Fisher Scientific 156499) at 5×10^6^ cells per flask per replicate. 24hrs post re-plating, cells were treated with 10x solutions (vehicle treatment or 1.6mM FeSO_4_ + 10uM RSL3) for either 8hr or 24hr. Post-treatment, cells were washed 2x with RT PBS and media replaced with complete microglia media. 2 days post-treatment the vehicle control flasks had DNA from 1×10^8^ cells per replicate isolated with Nucleobond CB 500 kit (Takara Bio 740509) according to manufacturer’s instructions. The 8hr and 24 hr ferroptosis-treated cultures were allowed to re-grow for 4-10 days. Once sufficiently re-grown, DNA from 1×10^8^ cells were isolated with Nucleobond CB 500 kit. DNA was stored at −80°C.

### CRISPR screen analysis

The sgRNA sequences were amplified using Guide-it™ CRISPR Genome-Wide Library PCR Kit (Takara, 632651) and subjected to the high-throughput amplicon sequencing on NextSeq500. 20bp of sgRNA sequences were first extracted using Cutadapt. The sgRNA counting and hit generation were done in MAGeCK and the downstream analysis were performed by MAGeCKFlute. The PCA plots were generated using edgeR and Glimma. Hits from the positive selection with a p value < 0.05 from each condition were imported into and intersected in R studio. PANTHER was used for the GO enrichment analysis. Each condition has two biological replicates.

### Screen of commercial ferroptosis-related compounds

The human microglia cell line was plated at 1,000 cells/ well of a 384-well plate. 24 hours post-plating the cells were co-treated with the commercial ferroptosis compound library (Selleckchem L6400) at 10μM or 2μM, depending on the stock concentration of 10mM or 2mM, and 1600μM FeSO_4_ + 1μM RSL3. 10μM for 535 of the compounds and 2μM for 11 of the compounds. 24 hours post-treatment, cell survivability was determined by cell titer glo (Promega G9241). Compounds were tested in triplicate.

### Quantification and statistical analysis

All quantification and statistical analyses were completed as described in the figure legends and methods. In brief, tri-culture biological replicates were considered independently differentiated cultures each with one to three technical replicates as individual wells. Statistical analysis across conditions was measured using one- or two-way analysis of variance (ANOVA). Dunnett’s, Tukey’s, or Sidak’s post hoc test was used as appropriate as indicated in figure legends. For gene expression change in scRNAseq or single Nucseq log-fold change cutoffs were >1.2 log fold and >.1 log2fold respectively with adjusted p value <0.05, as described in figure legends and methods. For the genome-wide CRISPR screen, two independent viral pools were used across two replicate cell cultures for each viral pool. For all experiments involving the SEC24B KO HAP1 line and its isogenic control, each biological replicate was considered and as an independent culture. Unpaired two-tail t test and one-way ANOVA with a Dunnett post hoc was used as indicated in the figure legends. The compound screen was performed with three technical replicates. Data for tri-culture death kinetics, FTH expression, and cell counts were log transformed to account for baseline shifts. For all panels where statistical significance is indicated, *p<0.05, **p<0.01, **p<0.001, and ****p<0.0001. Bar graphs display individual data points and report the data as the means ± SEM.

## Acknowledgements

AMP PD Differential expression analysis: Meaghan Cogswell, Dongyu Liu, Katherine W. Klinger, Stephen L. Madden, S. Pablo Sardi, Dinesh Kumar

AMP PD: Data used in the preparation of this article were obtained from the AMP PD Knowledge Platform. For up-to-date information on the study, https://www.amp-pd.org. “AMP PD – a public-private partnership – is managed by the FNIH and funded by Celgene, GSK, the Michael J. Fox Foundation for Parkinson’s Research, the National Institute of Neurological Disorders and Stroke, Pfizer, Sanofi, and Verily.

AMP PD Cohort: Clinical data and biosamples used in preparation of this article were obtained from the Parkinson’s Progression Markers Initiative (PPMI), and the Parkinson’s Disease Biomarkers Program (PDBP).

PPMI – a public-private partnership – is funded by the Michael J. Fox Foundation for Parkinson’s Research and funding partners, including [list the full names of all of the PPMI funding partners found at www.ppmi-info.org/fundingpartners]. The PPMI Investigators have not participated in reviewing the data analysis or content of the manuscript. For up-to date information on the study, visit www.ppmi-info.org.

Parkinson’s Disease Biomarker Program (PDBP) consortium is supported by the National Institute of Neurological Disorders and Stroke (NINDS) at the National Institutes of Health. A full list of PDBP investigators can be found at https://pdbp.ninds.nih.gov/policy. The PDBP Investigators have not participated in reviewing the data analysis or content of the manuscript.

For the Target ALS data, we would like to thank: The Target ALS Human Postmortem Tissue Core, New York Genome Center for Genomics of Neurodegenerative Disease, Amyotrophic Lateral Sclerosis Association and TOW Foundation, Discovery Bioinformatics Services, QIAGEN Digital Insights (QIAGEN Discovery Bioinformatics Services) especially Tiantian Ren and Hrisavgi Mangam. The Sanofi Translational Sciences Team especially John Green, Dominic Cammarata, Cheng Zhu, Mindy Zhang, Michael Dufault, Yi-Chien Chang, Lin An, Deepak Rajpal, and Katherine Klinger from Sanofi Translational Sciences.

## Author Contributions

Conceptualization: SKR, TRH

Methodology: SKR, TRH, DO

Investigation: SKR, TRH, YH, ET, LC, MS, SS, LG, CL, FP, EHJ, DK, MZ, JG, BZ, JP, JS, JCD, DR, DO

Visualization: SKR, TRH, ET, EHJ, JP

Project administration: TRH, DO

Supervision: TRH, DO

Writing – original draft: SKR

Writing – review & editing: SKR. TRH, DO, MZ, ET, LC, EHJ

## Data and materials availability

All data associated with this study are present in the paper or Supplementary Materials. AMP-PD data is available through the Terra platform by request and requires approval for access through the AMP-PD data use agreement. RNA-seq data will be deposited in GEO Accession.

