## Supplemental Material for "Microglia ferroptosis is prevalent in neurodegenerative disease and regulated by SEC24B"

**Supplementary Materials**

**Supplementary materials and methods**

**FACS on tri-culture**

Cells were washed 2x with PBS. Cells were then treated with 0.25% Trypsin + EDTA (Sigma T4049 for 6min in 37°C 5% $CO_2$ incubator. 1 volume of PBS + 2% FBS (Gibco A38400-01) was added to quench. Cells were combined from two wells and put through 40um cell strainers (BD Falcon 352235). Strained cell suspension was placed in a 1.7ml eppendorf tube on ice. Cell suspensions were spun at 4°C for 5min at 1500RPM. Supernatant was removed and cells were resuspended. Collected cells were resuspended in flow cytometry staining buffer (BD 554656) containing anti-GLAST (Miltenyi 130-118-344) PE-conjugated antibody for 10 minutes at room temperature. Samples were washed once, resuspended, and analyzed on a BD LSRII instrument. Results were analyzed in FlowJo software (Treestar).

**Immunocytochemistry**

Post treatment, supernatant was collected, and cells were washed 2x with RT PBS. Cells were fixed with 4% PFA (Fisher Scientific 50980495) for 15min at RT. Cells were washed 3x with PBS before being stored in PBS at 4°C. Cells were blocked with 5% BSA (Sigma A7030) + 0.1% Triton-X 100 (Sigma T9284) in PBS for 1hr at RT. Primary and secondary antibodies were prepared in blocking buffer. Cells were incubated in primary antibodies overnight at 4°C. Cells were washed 4x for 5min with PBS + 0.1% Tween20 (Sigma P9416). Cells were incubated in secondary antibodies for 1hr at RT in the dark. DAPI (Life Technologies D3571) was added during secondary incubation. Cells were washed 3x for 5min with PBS + 0.1% Tween20 and 1x in PBS. Cells were imaged and analyzed on Opera Phoenix.

Antibodies used for immunocytochemistry: mouse anti-ferritin (RnD Systems MAB93541, 1:500); rabbit anti-IBA1 (Wako 019-19741, 1:500); chicken anti-MAP2 (abcam ab92434, 1:1000); chicken anti-GFAP (Millipore AB5541, 1:500); donkey anti-mouse (invitrogen A21202, 1:500); donkey anti-rabbit (invitrogen A31572, 1:500); goat anti-chicken (invitrogen A21449, 1:500).

**Human immortalized microglia cell culture**

Human immortalized microglia (abm T3451) were cultured in complete microglia media: (DMEM Millipore D6546) +10% Tet-free FBS (Takara bio 631101) + 1x L-glutamine (Millipore #TMS-002-C) + 1x pen/strep (Millipore #TMS-AB2-C).

**Ferroptosis induction in human immortalized microglia**

The human microglia cell line (abm T3451) was plated at $5x10^3$ cells/ well of a 96 well plate. 24hrs post plating, cells were treated. Treatments included: 400uM $FeSO_4$, 800uM $FeSO_4$, and 1600uM $FeSO_4$, 1uM RSL3, 1uM $Ferrop_{Inh1}$, and 100nM $Ferrop_{Inh2}$. Cells were treated for 2,4, or 24 hours depending on the experiment. Cell viability was analyzed by Cell titer glo (Promega G9241).

**Lipidomics cell preparation**

The human microglia cell line was plated at 5x10$^6$ cells / T75 plate (Thermo Fisher Scientific 156499) in complete microglia media: (DMEM Millipore D6546) +10% Tet-free FBS (Takara bio 631101) + 1x L-glutamine (Millipore #TMS-002-C) + 1x pen/strep (Millipore #TMS-AB2-C). 24 hours later, cells were treated with 10x solutions. 10x solutions were: 1:500 DMSO, 1.6mM $FeSO_4$ + 1:500 DMSO, 1.6mM $FeSO_4$ + 10uM RSL3, 1.6mM $FeSO_4$ + 10uM RSL3 + 1uM $Ferrop_{Inh2}$, 1.6mM $FeSO_4$ + 10uM RSL3 + 10uM $Ferrop_{Inh1}$, 10uM RSL3, and 1uM $Ferrop_{Inh2}$. 2hrs and 4hrs post treatment, cells were collected. Cells were washed 2x with RT PBS. Cells were then treated with 0.25% Trypsin + EDTA for 5 min in 37°C incubator. Cells were quenched with 1 volume of complete microglia media. Cells were spun down at 4°C at 1200RPM for 5 min. Cells were washed and re-spun 2x with ice cold PBS. 1 million cells per sample were pelleted, supernatant was aspirated, flash frozen, and stored at -80°C before being sent for lipidomics analysis.

Lipidomic extraction solution preparation: 50 mL methanol was mixed with 50 mL chloroform. 0.2 mL acetic acid was added. 12(S)-HETE-d8 internal standard was added to get a final concentration of 10ng/mL. 15:0-18:1-d7-PE was added to get a final concentration of 20ng/mL.

**Unhydrolyzed sample preparation for lipidomics**

Cell pellet of 1x10$^6$ cells was resuspended with 0.2 mL ice cold water and resuspended by pipetting gently. The sample was transferred to a glass tube (with cap. 11 mL, 16 mm x 100 mm). 2 mL of extraction solution and 0.8 mL of water was added. Samples were vortexed (VWR, VX-2500 Multi-tube vortexer) for 5 min at the speed of 10 (max speed). Sample were incubated in the cold room for 20 min. Tubes were vortexed and then centrifuged for 10 minutes at 3000 rcf on Eppendorf 5810R centrifuge. The bottom layer was transferred to a new glass tube by using glass pasteur pipette and the organic solvent was dried by nitrogen. The dried lipid was resuspended in 100 µL methanol. Samples were transferred to LC-MS sample vials.

**Hydrolyzed sample preparation for lipidomics**

Cell pellet of 1x10$^6$ cells was resuspended with 0.2 mL ice cold water and resuspended by pipetting gently. The sample was transferred to a glass tube (with cap. 11 mL, 16 mm x 100 mm). 2 mL of extraction solution and 0.8 mL of water was added. Samples were vortexed (VWR, VX-2500 Multi-tube vortexer) for 5 min at the speed of 10 (max speed). Sample were incubated in the cold room for 20 min. Tubes were vortexed and then centrifuged for 10 minutes at 3000 rcf on Eppendorf 5810R centrifuge. The bottom layer was transferred to a new glass tube by using glass pasteur pipette. 50 ul 10 M KOH solution was added, mixed, and incubated at 37 °C for 2 h. 0.6 ml water was added, vortexed 2-5 min, and incubated in cold room for 20 min. Tubes were vortexed and then centrifuged for 10 min. The bottom layer was transferred to a new glass tube by using a glass pasteur pipette. The organic solvent was dried by nitrogen. The dried lipid was resuspended in 100 µL methanol. The sample was transferred to an LC-MS sample vial.

**H(p)ETE analysis by LC-MS**

LC separation was performed with a ACQUITY UPLC system. The multiple-reaction-monitoring (MRM) spectra were obtained with a Sciex Triple Quad 6500 mass spectrometer. A Waters Premier Acquity BEH C18 column (2.1 mm × 150 mm, 1.7 µm) was used for the LC separation.

The mobile phase consisted of (A) water/acetonitrile 75/25 with 10 mM ammonium acetate and (B) isopropanol/acetonitrile 50/50. The flow rate was 0.4 mL min–1. The solvent-gradient elution was changed as follows: 0 min, 0% B; 0.5 min, 0% B; 6 min, 50% B; 6.1 min, 100% B; 8 min, 100% B; 8.1 min, 0% B; 10 min, 0% B. Five microliter was injected. All data were acquired in negative-ion mode.

**H(p)ETE-PE analysis by LC-MS**

LC separation was performed with a ACQUITY UPLC system. The selected-ion-monitoring (SIM) spectra were obtained with Q-exactive HF high resolution mass spectrometer. Waters Acquity BEH HILIC column (1.7 um, 2.1 x 150 mm) was used for the LC separation. The mobile phase consisted of (A) 96% ACN, 2% MeOH, 1% Acetic acid, 1% $H_2O$, 5 mM Ammonium acetate and (B) 98% MeOH, 1% Acetic acid, 1% $H_2O$, 5 mM Ammonium acetate. The flow rate was 0.3 mL/min. The solvent-gradient elution was changed as follows: 0 min, 5% B; 2 min, 5% B; 15 min, 50% B; 16 min, 50% B; 16.1 min, 50% B; 25 min, 5% B. Five microliter were injected into the UPLC system. All data were acquired in negative-ion mode.

Standard curves were made with HETE-PE(C38:4) and HpETE-PE (C38:4) following the same extraction/sample prep.

**PE Profiling by High Resolution MS**

LC conditions were same as H(p)ETE-PE analysis. The profiling data (700-950Da in negative ion mode) were collected on Q-exactive HF high resolution mass spectrometer. PE class are eluted at a characteristic retention time on HILIC column (Waters Acquity BEH HILIC column 1.7 um, 2.1 x 150 mm) and PE identity are assigned by retention time and exact mass (<5ppm).

Profiling data is processed (aligned, peak picked, and clustered) by Expressionist 14.0 (Genedata, MS refiner). And statistical analysis (P-value and Fold change) is performed in Analyst (Expressionist 14.0, Genedata).

**PD blood bulk RNAseq analysis**

Data Availability: Unified AMP-PD Cohorts data is restricted and requires authorization for access: https://amp-pd.org/. Information and access to the Terra computing platform can be found at https://terra.bio/. Terra is developed by the Broad Institute of MIT and Harvard in collaboration with Verily Life Sciences. The Terra system was used for access to AMP-PD data and for computing. AMP-PD is a public-private partnership between the United States National Institutes of Health, non-profit organizations, and pharmaceutical and life sciences companies which aims to identify and validate new, impactful biological targets for PD therapeutics. For this analysis, PD Case and Control subjects enrolled in two large cohorts of the Accelerating Medicines Partnership - Parkinson's Disease (AMP-PD) Consortia were included: Parkinson’s Disease Biomarker Program (PDBP; 780 PD Cases, 504 Controls) and the Parkinson’s Progression Markers Initiative (PPMI; 816 PD cases, 617 Controls) (Fig. 5A). Although both PPMI and PDBP are case-control cohorts whose data are included in AMP-PD, it should be noted that these cohorts differ with respect to their inclusion criteria: PPMI inclusion criteria restricts enrollment to only those PD patients who have been diagnosed 2 years or less, whose baseline Hoehn & Yahr stage is I or II, and who are not expected to require medication for at least 6 months [1]. In contrast, PDBP enrolls PD patients across Hoehn & Yahr Stages, thus baseline patient data from PDBP includes a range of progression stages [2].

Data management, preprocessing and differential expression analysis were performed in Google Terra workspaces using R 3.6 and Python 3.7. Baseline differential gene expression analyses were performed for peripheral blood RNA sequencing data obtained at the point of study enrollment (baseline; BL) using the DESeq2 package [3]. The pathway analysis for DE gene sets was performed using Ingenuity Pathway Analysis software using adjusted p-value cutoff for pathway enriched <0.05 [4]. Transcriptomic quality control for samples was performed by the AMP-PD consortia. The decision tree for transcriptomics quality control is available on www.amp-pd.org. In this analysis, we have used the raw counts' output of Salmon v0.11.3 (PMCID: PMC5600148) for the abundant transcript estimation. Samples that were identified to have less than 50M reads, and outliers from the basic principal components analysis were flagged by the AMP-PD consortia and were not included in our further analysis. The DESeq2 R package (version 1.26.0) was used to identify differentially expressed genes among PD cases versus controls. DESeq2 is a method for differential expression analysis which may be applied to RNA-seq read counts per gene to estimate fold changes across conditions, in this case differences in gene expression measured from peripheral blood samples obtained from PD case and normal control subjects at point of study enrollment. As this is an exploratory analysis and validation, we have performed the analysis for the males and females study cohorts separately to identify potential differences between them. The design formula used for each differential expression analysis included age group as a covariate, sequencing plate number as batch variable, and diagnosis for the case vs. control comparison. Pre-processing and normalization of all RNAseq samples were done separately for each analysis for all expressed genes. Sample normalization and statistical testing for against a null hypothesis of no difference in gene expression between PD case and control subjects was performed using the Wald test, with p-values adjusted for multiple testing using the Benjamin and Hochberg method. Adjusted p-values of 0.05 and normalized gene log fold change in expression between conditions were then used as cutoffs for identification of protein-coding genes for use in downstream pathway analyses.

**Single nucleus RNASeq microglia analysis**

Control (n=3; 21,418 nuclei) and PD (n=3; 32,301 nuclei) snRNA samples data, sequencing, and analyses been previously reported [5, 6]. For initial broad types clustering, count matrices with nucleus barcodes and gene labels were loaded with R version 3.6.1/RStudio for sample integration and unsupervised clustering using Seurat Package version 3.1. For Quality Control (QC), nuclei were filtered following standard protocols based on examination of violin plots. Cutoffs 200 < nFeature_RNA < 9000 and percent.mt < 5 were used. Filtered matrices were then individually log-normalized by sample according to standard Seurat workflowsSample integration was performed in Seurat using the FindIntegrationAnchors and IntegrateData functions for 2000 variable features. Following integration and scaling according to Seurat package workflows, a range of clustering resolution values were trialed prior to broad cell type annotation (selected parameters: 2000 variable features, nPC = 20, and resolution = 0.8). Cluster-level expression of major cell type markers was examined and used to annotate cells contained within each cluster and confirm broad type assignments (Fig. 4).

Microglia Re-Clustering and Profiling: Nuclei identified as microglia by canonical marker expression were then subsetted from the Seurat object by cluster name ("Microglia") (Control, n = 849 nuclei; PD, n = 2274 nuclei) and this subset Seurat object was re-clustered using clustering parameters: nPCs = 5, nfeatures = 1000 and resolution = 0.2. All clusters generated contained nuclei from Control and PD samples. Cluster proportions and Dotplots for marker gene expression of interest are shown (Fig. 4).

Weighted Gene Co-expression Network Analysis (WGCNA): WGCNA was performed in R using the WGCNA package [7]. WGCNA was applied to the gene-gene co-expression matrix generated from the normalized cell-gene arrays from the integrated Seurat object containing microglia nuclei using soft thresholding power = 2 and deepSplit = 2. For each gene co-expression module identified by WGCNA, we then calculated the Pearson correlation between the module eigenvector and the dummy-encoded disease trait (PD or Control) in order to assess if any of these gene coexpression modules might be significantly correlated with disease state. Module genes co-expressed more actively in nuclei from a significantly correlated trait group are those which are positively correlated; negative module correlation suggests less active co-expression by trait.

**TargetALS**

Raw fastq RNA-Seq data files (1072 total) were provided by the NYGC ALS Consortium (Target ALS Release, June 2020) (https://www.targetals.org/research/resources-for-scientists/resource-genomic-data-sets/). These fastq files were generated using human postmortem tissue samples from the Target ALS postmortem tissue core and processed for RNA-Seq as previously described [8]. The raw fastq files were processed by Qiagen in OmicSoft ArrayStudio RNA-Seq analysis pipeline (version 10.1.1.3). In brief, the fastq files had quality control performed and then aligned to the Genome Reference Consortium Human Build 37 (GRCh37, GPL16791) using the proprietary OmicSoft Aligner [9]. After alignment, the gene level RPKM/FPKM/counts were determined using the EM algorithm in ArrayStudio as described previously [10]. Finally, pairwise differentially expressed genes were calculated using the DESeq2 v1.10.1 [3] in ArrayStudio comparing tissue-specific ALS patients vs. non-neurological controls. ALS patients with multiple neurological conditions were excluded from pairwise analysis. Heatmaps were generated using Excel. Differentially expressed genes (DEGs) were considered significant with a padj $< 0.05$.

**Protein Isolation**

*SEC24B* KO and isogenic control Hap1 cell lines (Horizon Discovery HZGHC001222c002 & C631) were plated on uncoated 6-well plates (Corning 3516) at $2x10^6$ cells per well. 24hrs post-plating, cells were washed 2x with RT PBS. Cells were then treated with 0.25% Trypsin + EDTA for 5 min in 37°C incubator. Cells were quenched with 1 volume of HAP1 media (Iscove's Modified Dulbecco's Medium (IMDM) (Gibco 12440-053) + 20% FBS (Gibco A38400-01)). Cells were spun down at RT at 1200RPM for 5 min. Supernatant was aspirated and cells were resuspended in PBS and spun down at RT at 1200RPM for 5 min. Supernatant was aspirated and cell pellet was resuspended in 100uL 1x RIPA buffer (Boston BioProducts BP-115) with cOmplete protease inhibitor (Roche 11697498001). Cells were briefly vortexed and left on ice for 10 min. Samples were spun at 14,000 RPM for 10 min at 4°C. Protein concentration was determined by BCA (Thermo Scientific 23227).

##### Western blot

Protein samples were prepared in LDS sample buffer (Life Technologies B0007) and reducing agent (Life Technologies B0009). Samples were boiled for 10min at 70°C and then run in 4-12% Bis-Tris gels (Invitrogen NP0321BOX) in 1x MOPS running buffer (Life technologies NP0001) at 150V for 1hr. Gel was dry transferred with iBlot2 (Invitrogen). Blots were blocked for 1hr at RT in Intercept Blocking Buffer (Li-Cor 927-60001). Blots were incubated in primary antibody solution (Intercept Blocking Buffer) with rabbit anti-SEC24B (Cell Signaling Technology 12042S, 1:1,000) and mouse anti-β-actin (Sigma A5441, 1:5,000) at 4°C overnight. Blots were incubated in Secondary antibody solution (Intercept Blocking Buffer + 0.1% Tween-20) with Donkey anti-Mouse 680 (Li-Cor 926-68072, 1:2,000) and Donkey anti-Rabbit 800 (Li-Cor 926-32213, 1:2,000) at RT for 1hr. Blots ere imaged on Odyssey CLx.

##### RNA isolation and qRT-PCR

RNA was isolated with RNeasy Plus Mini Kit (Qiagen 74134) according to manufacturer's instructions. cDNA was synthesized from RNA with SuperScript VILO (Invitrogen 11755-050). RNA expression was quantified with taqman probes (Thermo Fisher Scientific) for *SEC24B* (Hs00197035_m1) and *RPL37A* (Hs01102345_m1) and TaqMan Gene Expression Master Mix (Applied Biosystems 4369016). 30ng of cDNA was used per well, with three technical replicates per sample. qRT-PCR was run on Applied Biosystems QuantStudio 7 Flex. Expression levels were normalized to *RPL37A.*

#### Supplementary Figures

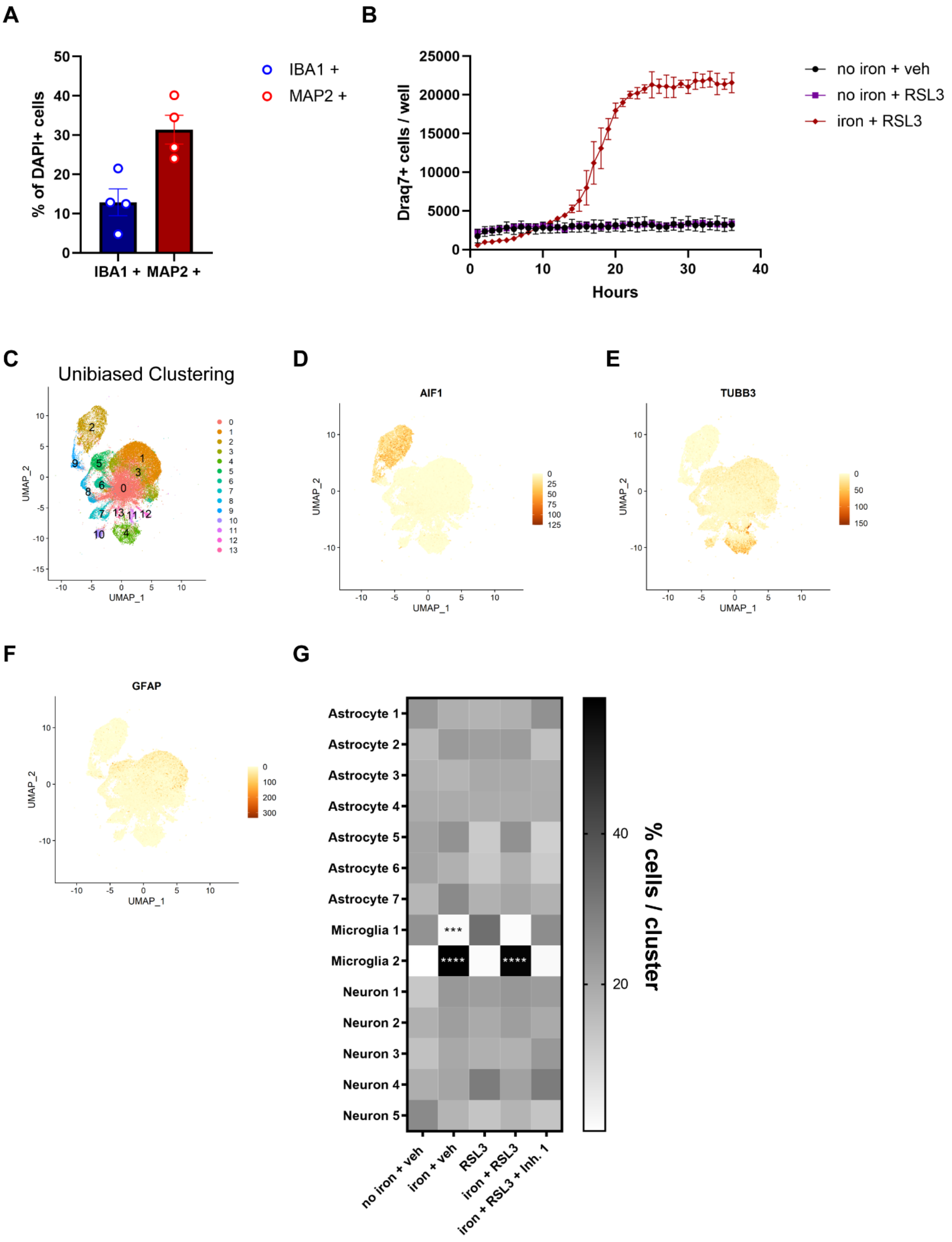

**Fig. S1. Tri-culture cell type validation and death kinetics of RSL3 only. (A)** Quantification of percent IBA1+ and MAP2+ cells by ICC in tri-culture (n=4). Error bars represent SEM. **(B)** Death kinetics of no iron + RSL3 compared to no iron + veh and iron + RSL3. (n=1, 3 technical replicates). Error bars represent SEM. **(C)** Unbiased clustering of scRNAseq for tri-cultures (n=3). **(D)** to **(F)** Cell type identification of gene expression by **(D)** *AIF1* for microglia, **(E)** *TUBB3* for neurons, and **(F)** *GFAP* for astrocytes. **(G)** Quantification of subclusters per condition. Two-way ANOVA, Tukey post hoc. ***$p<0.001$, ****$p<0.0001$.

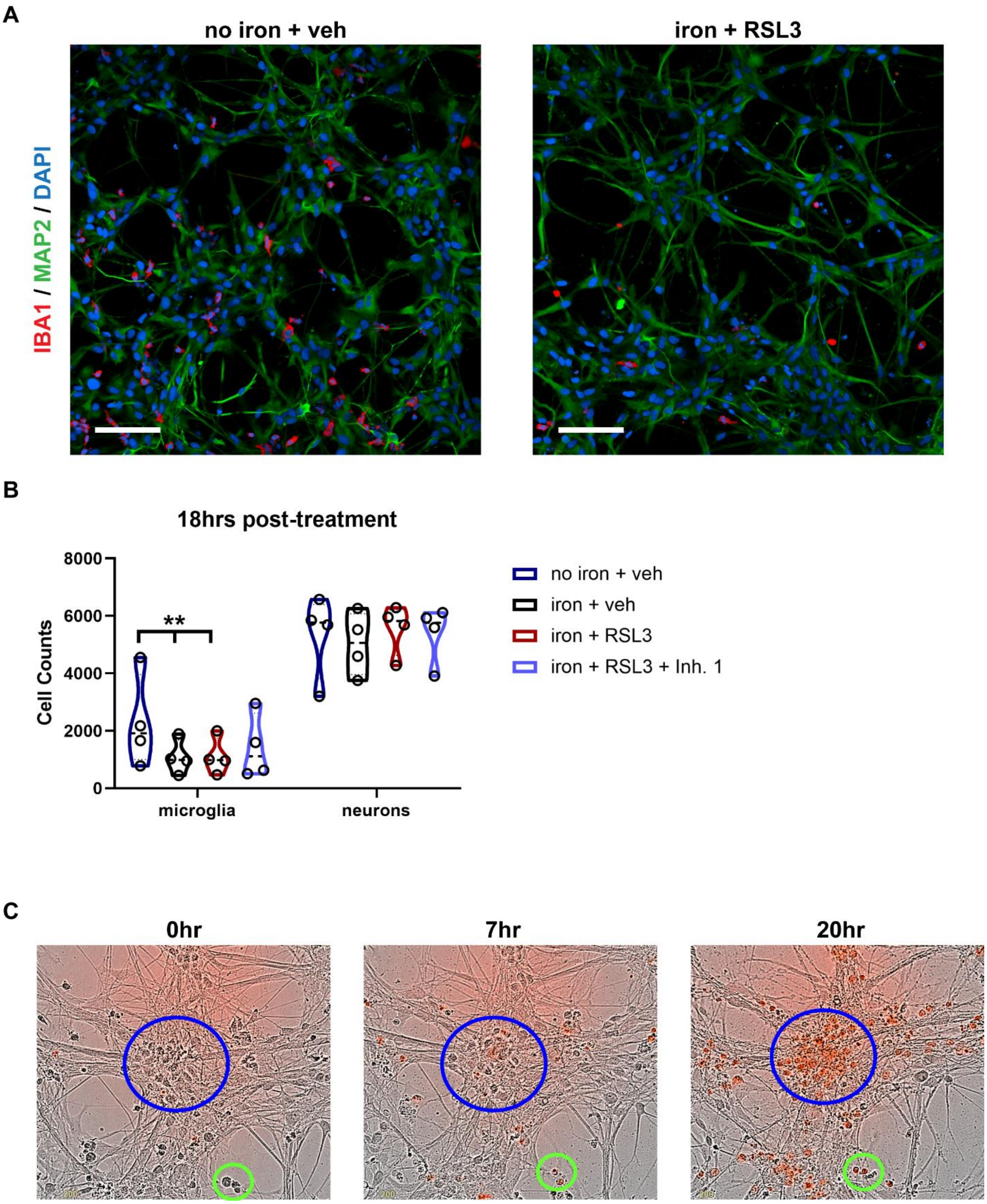


**Fig. S2. Ferroptosis is induced in microglia, followed by neurons. (A)** Representative images of IBA1+ microglia (red) and MAP2+ neurons (green) in tri-culture after 18hr exposure to no iron + veh or iron + RSL3 treatments. **(B)** Quantification of microglia and neurons 18hrs post-treatment (n=4). One-Way ANOVA, Dunnett post hoc, log transformed. **p<0.01. **(C)** Representative images of tri-culture treated with iron +RSL3 at 0hr, 7hrs, and 20hrs post-treatment. Blue circle identifies a neuron cluster. Green circle identifies microglia by morphology. Red cells are Draq7+ cells, indicating microglia death by 7hrs and neuronal death around 20hrs post-treatment.

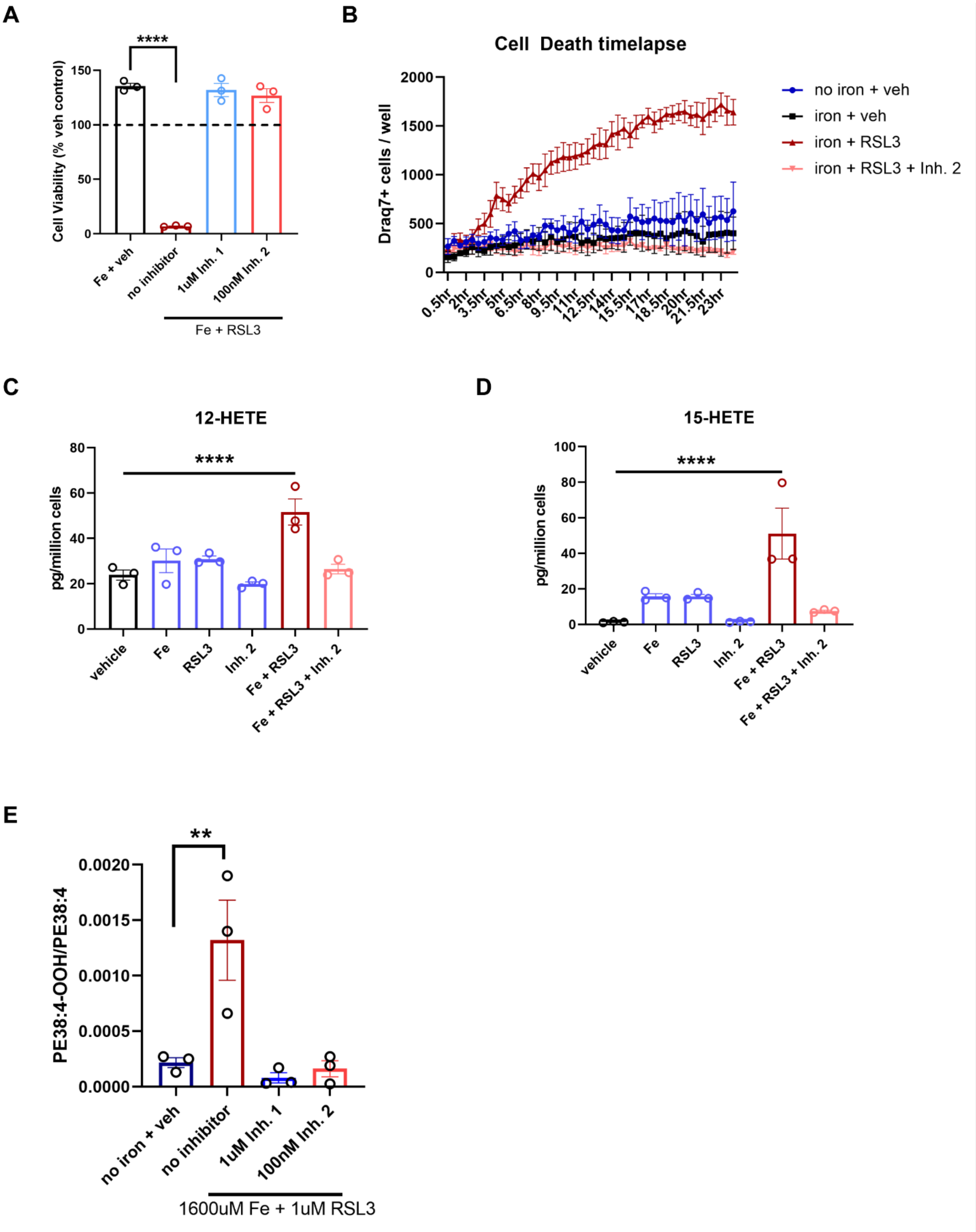


**Fig. S3. Lipidomic analysis reveals distinct ferroptotic signature in microglia (A)** Ferroptosis induction (800uM iron + 1uM RSL3) ± inhibitors (1uM Inh. 1 / 100nM Inh. 2) in human immortalized microglia cell line. (n=3). One-way ANOVA, Dunnett post hoc.

***p<0.001. Error bars represent SEM. **(B)** Draq7+ death kinetics of human immortalized microglia exposed to 400uM iron + 1uM RSl3 ± 100nM $\text{Ferrop}_{\text{Inh2}}$. (n=3 wells). Error bars represent SEM. **(C)** and **(D)** Lipidomic analysis shows increased free 12-HETE and 15-HETE in immortalized human microglia 2hr post-treatment. (n=3). One-way ANOVA, Sidak post hoc. ****p<0.0001. Error bars represent SEM. **(E)** Production of 1-SA-2-15-HpETE-PE ± or 1µM $\text{Ferrop}_{\text{Inh1}}$ or 100nM $\text{Ferrop}_{\text{Inh2}}$. (n=3). One-way ANOVA, Dunnett post hoc. **p<0.01. Error bars represent SEM.

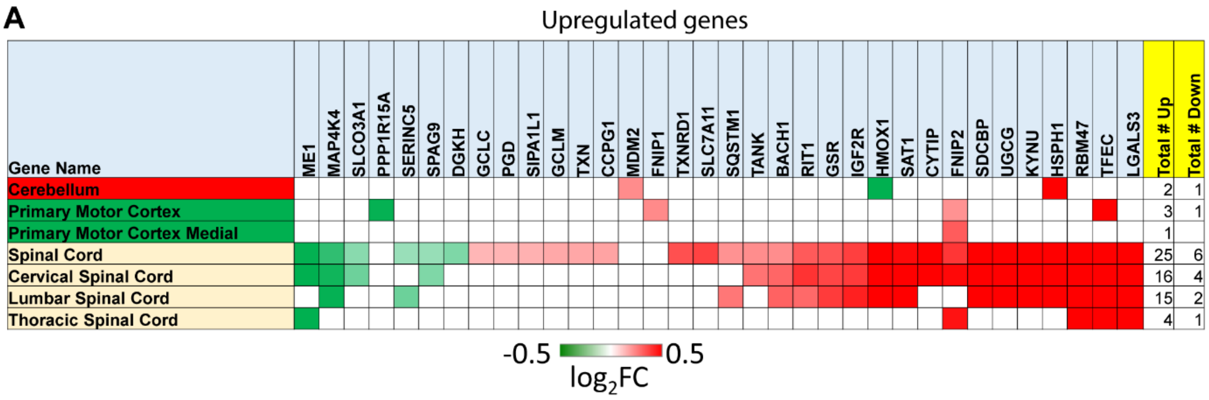


**Fig. S4. Ferroptosis-related genes are significantly upregulated in ALS patient spinal cord.** (**A**) Heatmap of upregulated genes in tri-culture ferroptotic microglia in ALS patients compared to control. Boxes shown in red or green are genes with $P_{adj}$<0.05. The number of samples, area, and disease state analyzed are reported in table S1.

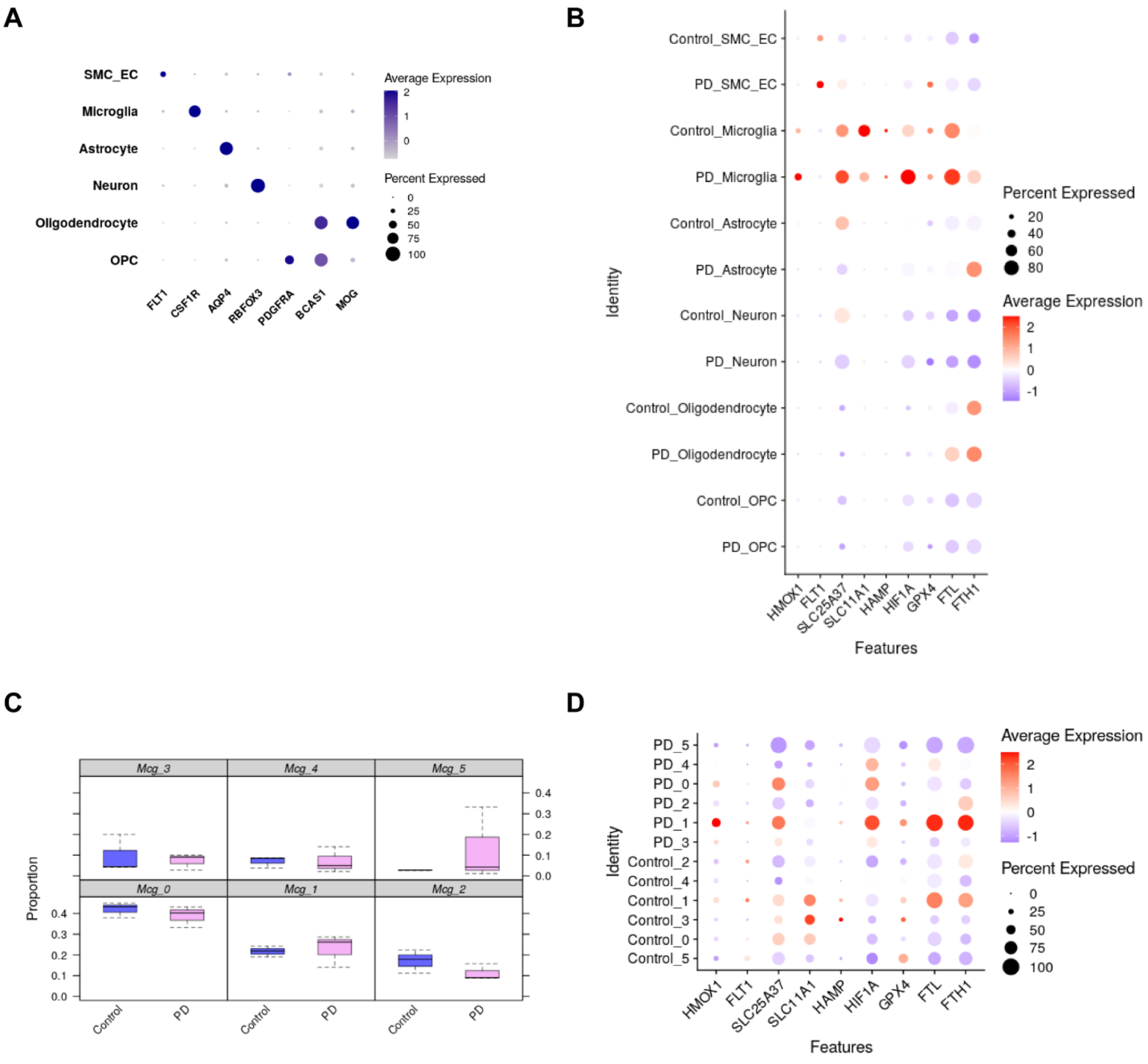


**Fig. S5. Cell type identification in single nucSeq and expression analysis of MS microglia iron-related signature in each cell type. (A)** Dot plot for cell-type specific gene expression for endothelial cells, microglia, astrocytes, neurons, oligodendrocytes, and OPCs. **(B)** MS microglia iron-related signature was upregulated specifically in PD microglia compared to control and other cell types. (C) Proportions for each of the six microglia subclusters across control and PD. **(D)** Re-clustering of microglia nuclei reveals subpopulation of PD microglia with greater expression of signature genes.

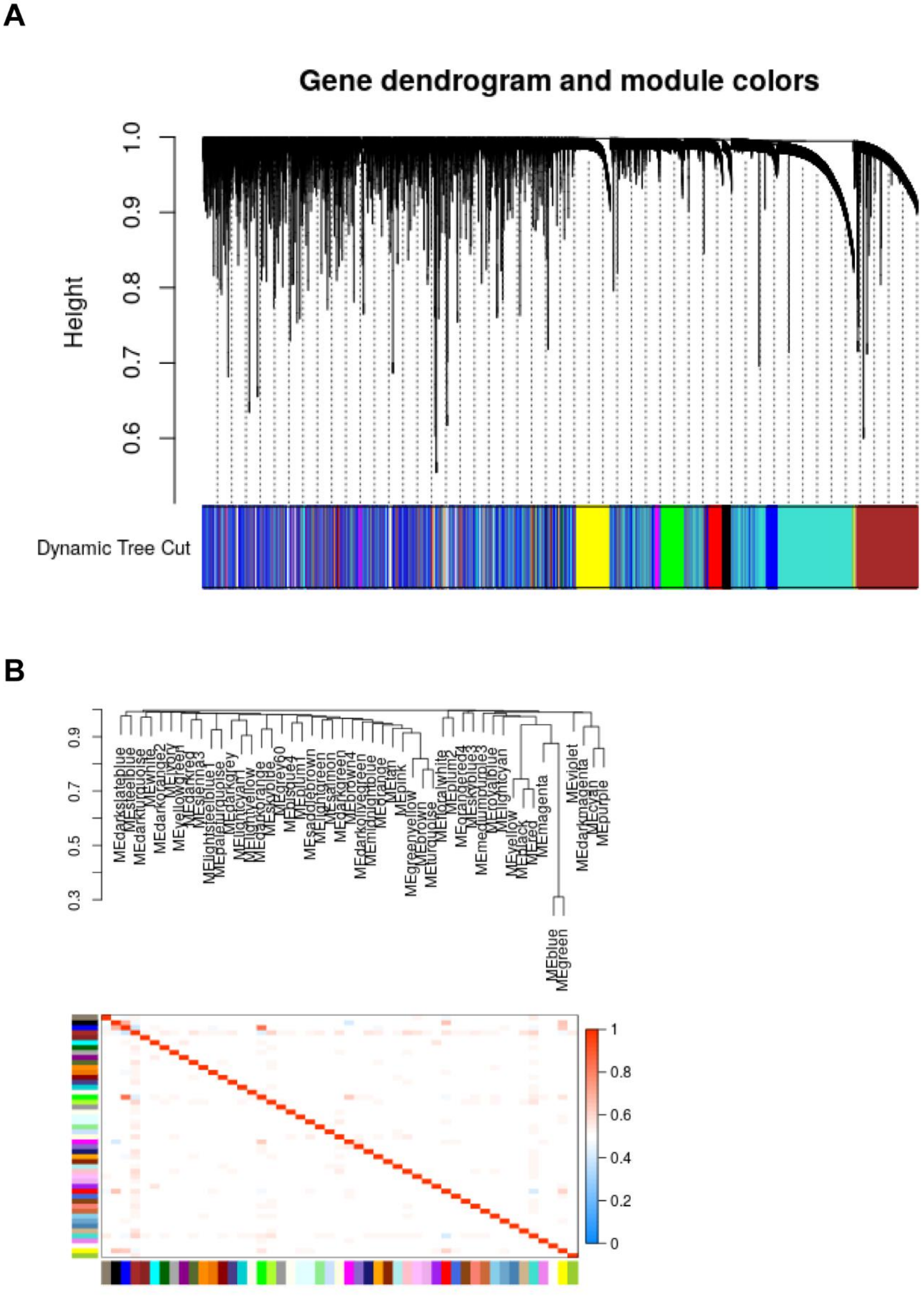
A
Gene dendrogram and module colors
Height
1.0
0.9
0.8
0.7
0.6
Dynamic Tree Cut
B
0.9
0.7
0.5
0.3
MEblue
MEgreen
1
0.8
0.6
0.4
0.2
0

**Fig. S6. Quality control of WGCNA. (A)** Dendogram of modules. **(B)** Eigengene network heatmap.

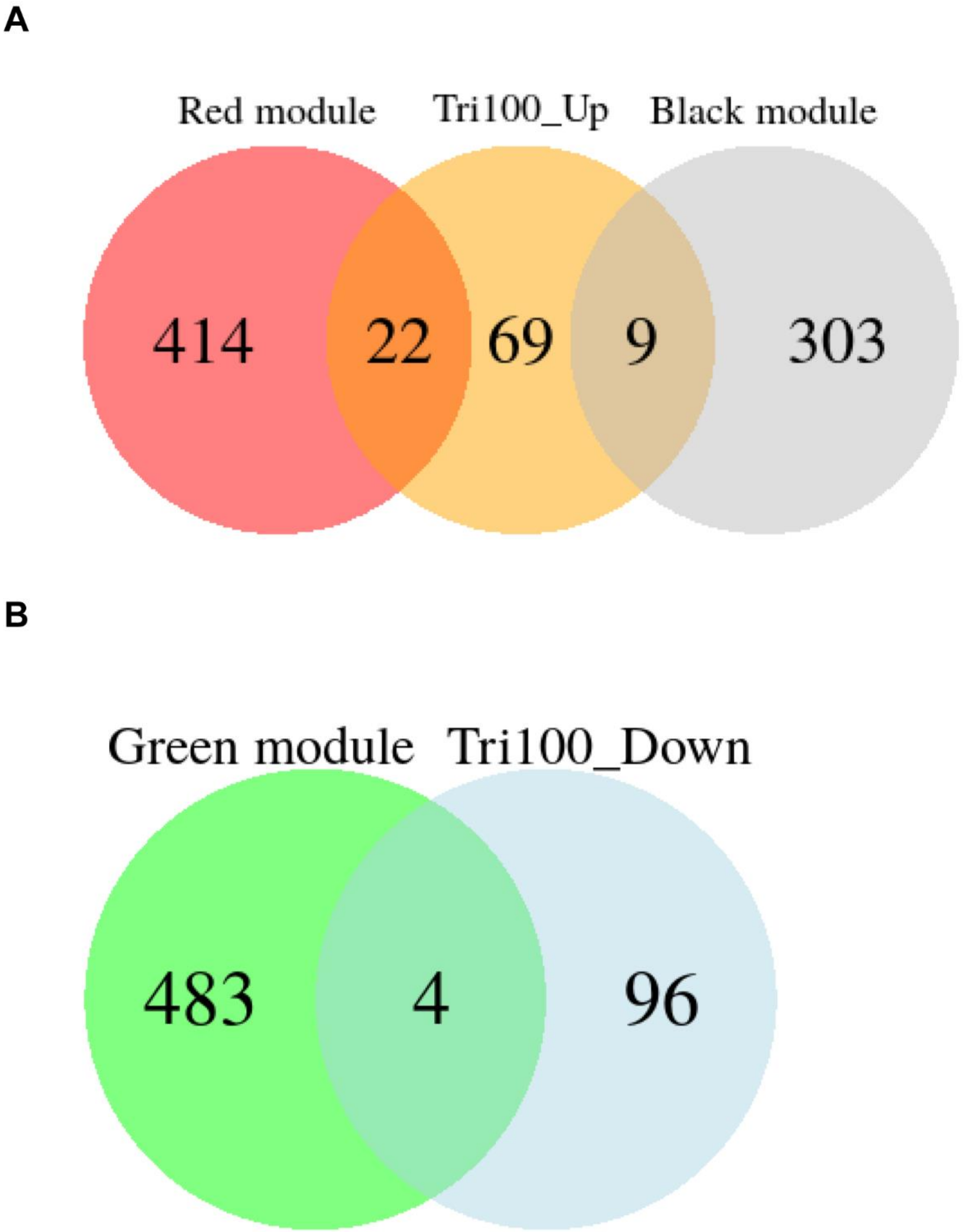


**Fig. S7. Venn diagram of microglia tri-culture ferroptosis signature and trait-relationship modules. (A)** Venn diagram of shared genes of the top 100 upregulated genes in the tri-culture ferroptosis signature and black and red modules. **(B)** Venn diagram of shared genes of the top 100 downregulated genes in the tri-culture ferroptosis signature and green module.

A

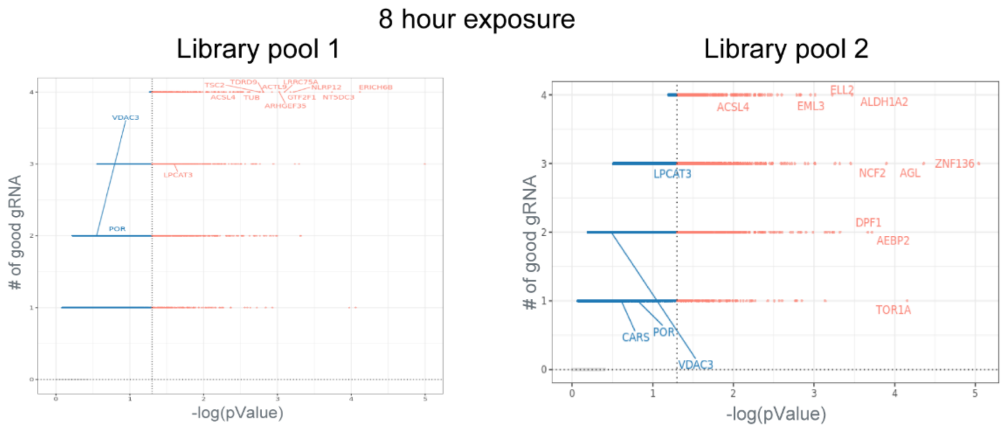
8 hour exposure
Library pool 1
Library pool 2
### of good gRNA
-log(pValue)
VDAC3
POR
LPCAT3
ELL2
ALDH1A2
ACSL4
EML3
ZNF136
NCF2
AGL
DPF1
AEBP2
TOR1A
CARS

B

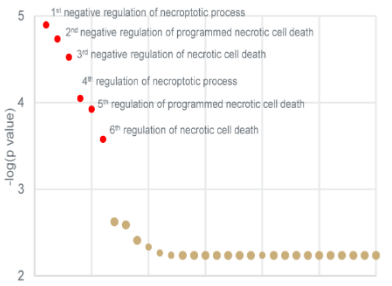
-log(p value)
1st negative regulation of necroptotic process
2nd negative regulation of programmed necrotic cell death
3rd negative regulation of necrotic cell death
4th regulation of necroptotic process
5th regulation of programmed necrotic cell death
6th regulation of necrotic cell death

C

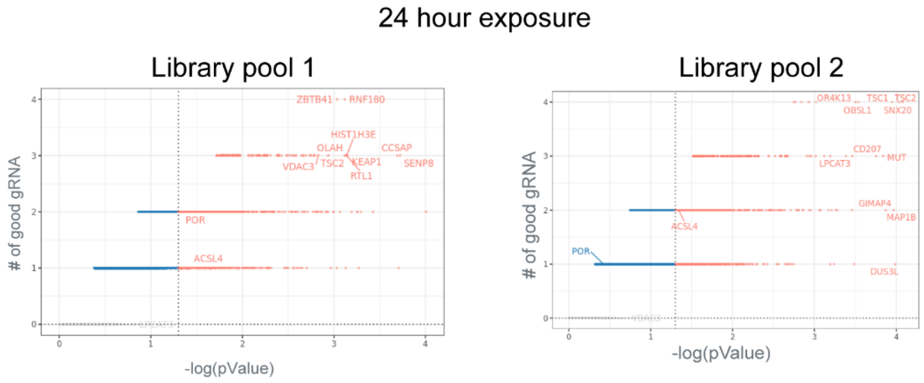
24 hour exposure
Library pool 1
Library pool 2
### of good gRNA
-log(pValue)
ZBTB41
RNF180
HIST1H3E
OLAH
CCSAP
VDAC3
TSC2
KEAP1
SENP8
RTL1
POR
ACSL4
OR4K13
TSC1
TSC2
OBSL1
SNX20
CD207
MUT
LPCAT3
GIMAP4
MAP1B
DUS3L

D

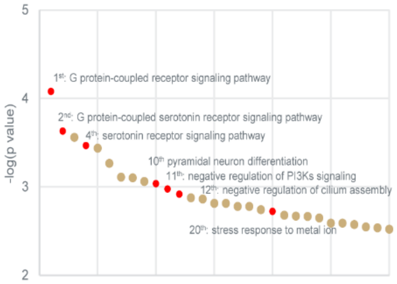
-log(p value)
1st: G protein-coupled receptor signaling pathway
2nd: G protein-coupled serotonin receptor signaling pathway
4th: serotonin receptor signaling pathway
10th pyramidal neuron differentiation
11th: negative regulation of PI3Ks signaling
12th: negative regulation of cilium assembly
20th: stress response to metal ion

**Fig. S8. Crispr Screen additional information. (A)** Log p value and number of gRNAs identified for each gene across both library pools at the 8-hour timepoint. **(B)** Top pathways associated with hits from the 8-hour exposure. **(C)** Log p value and number of gRNAs identified for each gene across both library pools at the 24-hour timepoint. **(D)** Top pathways associated with hits from the 24-hour exposure.

#### Supplementary Tables

##### Table S1. Target ALS sample numbers

| Table S1. Target ALS Tissue Breakdown | | | |
|---|---|---|---|
| **Tissues** | **ALS** | **Non-Neurological Control** | **Both** |
| cerebellum | 114 | 13 | 127 |
| frontal cortex | 95 | 13 | 108 |
| motor cortex lateral | 80 | 11 | 91 |
| motor cortex medial | 80 | 12 | 92 |
| motor cortex unspecified | 9 | 1 | 10 |
| motor cortex | 169 | 24 | 193 |
| occipital cortex | 46 | 6 | 52 |
| cervical spinal cord | 92 | 12 | 104 |
| lumbar spinal cord | 79 | 10 | 89 |
| thoracic spinal cord | 44 | 8 | 52 |
| spinal cord | 215 | 30 | 245 |

##### Table S2. Nuclei counts for each cell type in snRNAseq dataset

| | OPC | Oligodendrocyte | Neuron | Astrocyte | Microglia | SMC_EC | row_sum |
|---|---|---|---|---|---|---|---|
| | <int> | <int> | <int> | <int> | <int> | <int> | <dbl> |
| Control1 | 215 | 3789 | 375 | 362 | 215 | 4 | 4960 |
| Control3 | 464 | 3362 | 1534 | 1164 | 314 | 57 | 6895 |
| Control8 | 412 | 7660 | 1376 | 634 | 320 | 40 | 10442 |
| PD2 | 254 | 9193 | 125 | 621 | 524 | 9 | 10726 |
| PD4 | 252 | 4342 | 3643 | 2608 | 1020 | 147 | 12012 |
| PD9 | 259 | 6513 | 982 | 1259 | 470 | 80 | 9563 |

##### Table S3. PPMI and PDBP sample numbers

| Study | All | # Participants | | Gender | |
|---|---|---|---|---|---|
| | | Case | Control | Female | Male |
| PPMI | 1433 | 816 | 617 | 669 | 764 |
| PDBP | 1284 | 780 | 504 | 574 | 710 |

**Table S4. Dysregulated genes in each group associated with ferroptosis pathway**

| **Ferroptosis Pathway Genes Differentially Expressed in PD v, Control** | |
|---|---|
| **PPMI Female (n=10)** | ALOX15, GCLC, GPX4, H2BC10, H2BC15, H2BC17, H2BC5, H2BC8, MAPK3, RRAS2 |
| **PDBP Female (n = 19)** | ABCA1, ACSL4, ALOX15B, ALOX5, BAP1, BID, H2AC18/H2AC19, H2AZ1, H2BC12, HMGCR, HMOX1, MAP2K2, MAPK1, MAPK3, RAF1, RALB, SQSTM1, STAT3, TXNRD1 |
| **PPMI Male (n=7)** | ABCA1, ALOX5, H2BU1, MAPK3, SLC1A5, STAT3, TFRC |
| **PDBP Male (n=12)** | ABCA1, ACSL4, ALOX15, ALOX5, GCLC, H2AZ1, RALB, SAT1, SLC1A5, SLC7A11, SREBF2, TXNRD1 |

#### Table S5. Full hit list from ferroptosis compound screen

| compound name | percent recover | conc used | target | pathway |
|---|---|---|---|---|
| Xanthotoxol | 158.3764471 | 10uM DMSO | Others | 5-HT Receptor |
| BAY 87-2243 | 87.10295311 | 10uM DMSO | Inflammation/Immunology | Angiogenesis |
| Sesamol | 97.73483611 | 10uM DMSO | E3 Ligase ,p53 | antioxidant |
| Ethoxyquin | 75.584162 | 10uM DMSO | NF-κB,HDAC,Histone Acetyltransferase,Nrf2 | antioxidant |
| Diphenylamine Hydrochloride | 82.05480531 | 10uM water | Others | antioxidant |
| Lauryl gallate | 70.36480757 | 10uM DMSO | Others | antioxidant |
| (+)-Delta-Tocopherol | 77.59477972 | 10uM DMSO | iNOS | antioxidant |
| Octyl gallate | 74.85572818 | 10uM DMSO | NF-κB | antioxidant |
| Cyclic Pifithrin-α hydrobromide | 72.77337001 | 10uM DMSO | Cancer | Apoptosis |
| WNK463 | 70.42145001 | 10uM DMSO | Vitamin | Apoptosis |
| Tenovin-1 | 149.1821169 | 10uM DMSO | GMNN - geminin, DNA replication inhibitor (human) | Apoptosis |
| Necrostatin-1 | 83.85006199 | 10uM DMSO | ALK,c-Met | Apoptosis |
| 3,6'-Disinapoyl sucrose | 76.23339408 | 10uM DMSO | FXR receptor | Bcl-2 |
| Isosilybin | 73.41418741 | 10uM DMSO | HIF | Cancer |
| Schisandrin C | 95.48330333 | 10uM DMSO | Histamine Receptor | Cancer |
| Epiberberine | 89.4575386 | 10uM DMSO | TNF-alpha | DNA replication |
| Equol | 72.21767315 | 10uM DMSO | 5-HT Receptor | Endocrinology & Hormones |
| Psoralidin | 82.98420834 | 10uM DMSO | P450 (e.g. CYP17) | Endocrinology & Hormones |
| Curcumin | 129.6421671 | 10uM DMSO | Estrogen/progestogen Receptor | Epigenetics |
| Boldine | 87.34264647 | 10uM DMSO | Others | FXR receptor |
| Schisanhenol | 82.17574014 | 10uM DMSO | Others | glucuronosyltransferases |
| Olivetol | 71.0489686 | 10uM DMSO | Ferroptosis | GPCR & G Protein |
| Thymol | 77.85836852 | 10uM DMSO | NADPH-oxidase | Immunology & Inflammation |
| 2-Acetylphenothiazine (ML171) | 70.71460427 | 10uM DMSO | AChR | Immunology & Inflammation |
| Trolox | 93.4535961 | 10uM DMSO | Sodium Channel | Metabolism |
| Galangin | 83.13078825 | 10uM DMSO | Immunology & Inflammation related | Metabolism |
| ML355 | 75.13140889 | 10uM DMSO | Others | Metabolism |
| Ferrostatin-1 (Fer-1) | 80.03178369 | 10uM DMSO | 3,6'-Disinapoyl sucrose | Metabolism |
| GKT137831 | 79.43502429 | 10uM DMSO | Others | NADPH-Oxidase |
| Levocetirizine Dihydrochloride | 84.72345445 | 10uM DMSO | Lipoxygenase | Neuronal Signaling |
| Jatrorrhizine | 78.91407161 | 10uM DMSO | Others | Neuronal Signaling |
| 8-OH-DPAT (8-Hydroxy-DPAT) | 83.49121491 | 10uM DMSO | Others | Neuronal Signaling |
| JSH-23 | 96.08833966 | 10uM DMSO | CYP3A474µM | NF-κB |
| ATP | 111.0616153 | 10uM DMSO | p53 | Others |
| Rhapontigenin | 191.1995477 | 10uM DMSO | Estrogen/progestogen Receptor | proliferation |
| Ferulaldehyde | 75.0495201 | 10uM DMSO | Cannabinoid Receptor | proliferation |
| Crizotinib (PF-02341066) | 88.45740205 | 10uM DMSO | NADPH-oxidase | Protein Tyrosine Kinase |
| Zonisamide | 78.20997873 | 10uM DMSO | Serine/threonin kinase | Transmembrane Transporters |
| Demethoxycurcumin | 97.50980203 | 10uM DMSO | Others | Wnt/β-catenin |
